# The genomic landscape of swine influenza A viruses in Southeast Asia

**DOI:** 10.1101/2023.02.10.527943

**Authors:** Michael A Zeller, Jordan Ma, Foong Ying Wong, Sothrya Tum, Arata Hidano, Hannah Holt, Ty Chhay, Sorn San, Dina Koeut, Bunnary Seng, Sovanncheypo Chao, Giselle GK Ng, Zhuang Yan, Monidarin Chou, James W Rudge, Gavin JD Smith, Yvonne CF Su

## Abstract

Swine are a primary source for the emergence of pandemic influenza A viruses. The intensification of swine production, along with global trade, has amplified the transmission and zoonotic risk of swine influenza virus (swIAV). Effective surveillance is essential to uncover emerging virus strains, however gaps remain in our understanding of the swIAV genomic landscape in Southeast Asia. By collecting more than 4,000 nasal swabs and 4,000 sera from pigs in Cambodia, we unmasked the co-circulation of multiple lineages of genetically diverse swIAV of pandemic concern. Genomic analyses revealed a novel European avian-like H1N2 swine reassortant variant with North American triple reassortant internal genes, that emerged approximately seven years before its first detection in pigs in 2021. Using phylogeographic reconstruction, we identified south central China as the dominant source of swine viruses disseminated to other regions in China and Southeast Asia. We also identified nine distinct swIAV lineages in Cambodia, which diverged from their closest ancestors between two to 15 years ago, indicating significant undetected diversity in the region, including reverse zoonoses of human H1N1/2009 pandemic and H3N2 viruses. A similar period of cryptic circulation of swIAVs occurred in the decades before the H1N1/2009 pandemic. The hidden diversity of swIAV observed here further emphasizes the complex underlying evolutionary processes present in this region, reinforcing the importance of genomic surveillance at the human-swine interface for early warning of disease emergence to avoid future pandemics.

## Introduction

Global swine populations play an integral role in the emergence of zoonotic influenza and pandemic viruses by providing a suitable environment for reassortment and human adaptation of avian and mammalian influenza viruses [1-3]. As pork production has dramatically increased over the past 50 years (Fig. 1), international trade and movement have further amplified the zoonotic risks at the human-pig interface [4-6]. In 2020, Southeast Asian countries had estimated swine inventories of 1.9 million head in Cambodia, 4.3 million in Laos, 7.5 million in Thailand, and 22 million in Vietnam [6, 7]. The impact of intensified commercial swine production for the emergence and spread of swine influenza A virus (swIAV) is still unknown, particularly in Southeast Asia where epidemiological and genetic data on swIAV are sparse.

**Fig. 1.**
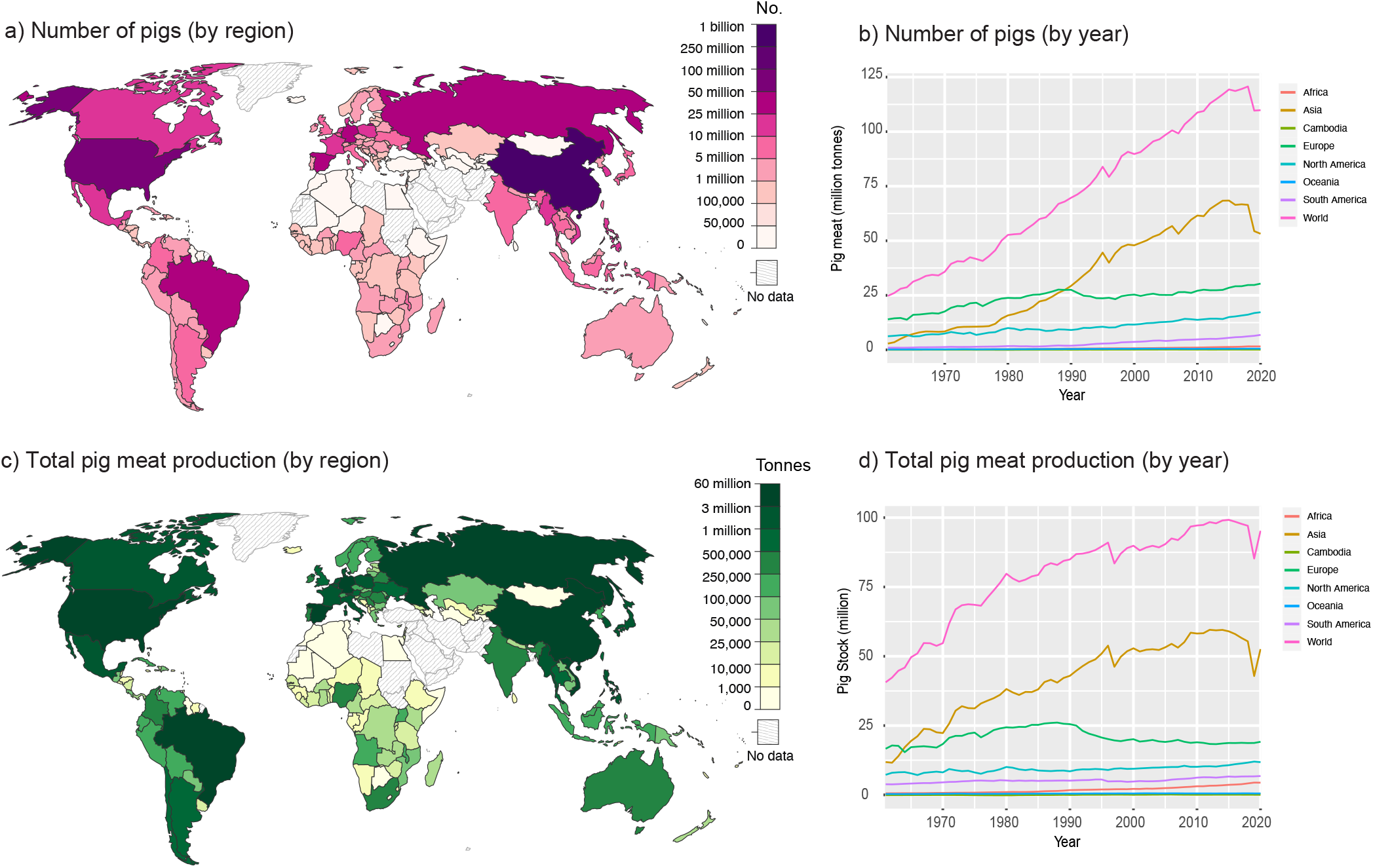
Comparison of pig production in different geographic regions. (a) Global distribution of pig populations in 2020. (b) Number of pigs in the world by year. (c) Global distribution of total pig meat production in 2020. (d) Total pig meat production by year. Source: The United Nations Food and Agriculture Organization (FAO) and Our World in Data (OWID).

Establishing the exact origin of the human H1N1/2009 pandemic (H1N1/pdm09) virus was prevented by the lack of available sequence data from swine [1, 2, 8, 9]. Dated phylogenies showed that the swIAV ancestors of the H1N1/pdm09 virus were unsampled for at least 10 years [8]. It was not until 2016 that reassortants of European avian-like (EA) and North American swine viruses found in Mexican swine were identified as H1N1/pdm09 precursors [10]. Analysis of swIAV viruses collected in Hong Kong from 1998–2010 showed that virus population dynamics were dominated by repeated intercontinental movement and reassortment of swIAVs that produced diverse virus populations [3]. Population based simulation of virus movements conducted in 2015 suggested that swIAV diversity from large pig populations in China and the United States, which are the source of most swIAV genomes, did not represent global diversity and that surveillance efforts should be extended to other regions [11]. These studies demonstrate the central role of global swine movements in facilitating the mixing of independently evolving swIAV populations that leads to reassortment and the generation of novel swIAV strains.

Multiple lineages of swIAV have been detected in pigs across Southeast Asia [12-18]. In Thailand, the H1N1/pdm09 virus was the prevailing subtype detected in swine from 2011 to 2014 [14], co-circulating with classical swine (CS) H1, EA H1N1, and H3N2 lineages [13]. Multiple H1 and H3 genotypes have been generated in swine from Thailand through frequent reassortment between these lineages [12, 13]. Surveillance studies from Vietnam between 2010–2013 detected H1N1, H1N2, and H3N2 subtype IAV in swine [15-17]. While in Myanmar, H1N1/pdm09, H1N2, and multiple H3 lineages have been detected from 2017– 2019 [18].

Influenza A viruses exhibit broad host tropism outside of the natural reservoir of wild waterfowl and can infect a wide range of species, including humans, swine, and domesticated fowl such as chicken, that have frequent human contact in live-bird markets throughout Asia [19]. There have been multiple documented cases of zoonotic [8, 20-23] and reverse-zoonotic [24-28] transmission and establishment of IAV in multiple hosts. Cambodian smallholders often keep swine alongside chicken and ducks [29], creating an environment favourable to cross-species transmission [30, 31]. However, there are very few swIAV sequences available from Cambodia and the rest of Southeast Asia, and the lack of systematic surveillance infrastructure has created gaps in our ability to identify emerging pathogens in pigs and their potential for zoonotic spillover [7].

In this study, we collected over 4,000 nasal swabs and 4,000 sera from pigs in Cambodia to reveal the prolonged undetected circulation and establishment of diverse swIAV lineages in Cambodia, including human H1N1/pdm09 and H3N2 viruses introduced to swine through reverse-zoonosis. We uncovered a novel EA H1N2 reassortant genotype generated through the intercontinental movement of different viral lineages, and phylogeographic reconstruction demonstrated that China is the leading source contributing to dissemination of EA swine viruses in Asia. Our results unmask an increasingly complex genomic landscape of swIAV in Southeast Asia that is shaped by repeated introduction and reassortment of new virus lineages, a process that is known to heighten pandemic risk.

## Results

### Detection and characterization of swine influenza A viruses in slaughterhouse pigs

From March 2020 to July 2022, swine influenza surveillance was conducted in 18 pig slaughterhouses in Cambodia. We collected a total of 4,089 nasal swabs and 4,069 sera samples from individual pigs from different districts in four neighboring provinces (Kampong Speu, Kandal, Phnom Penh, and Takeo) (Fig. 2). Of these, 72 sampled pigs (1.8%) were positive for influenza A virus by RT-qPCR (Table 1). Kandal province exhibited a higher positivity rate of IAV by RT-qPCR (4.5%) compared to the three other provinces (0.2–1.8%). All sera were tested for the presence of anti-nucleoprotein IAV antibodies by enzyme-linked immunosorbent assay (ELISA). The overall IAV seroprevalence rate was 30.0–36.9% in sampled pigs across the four provinces (Table 1).

**Fig. 2.**
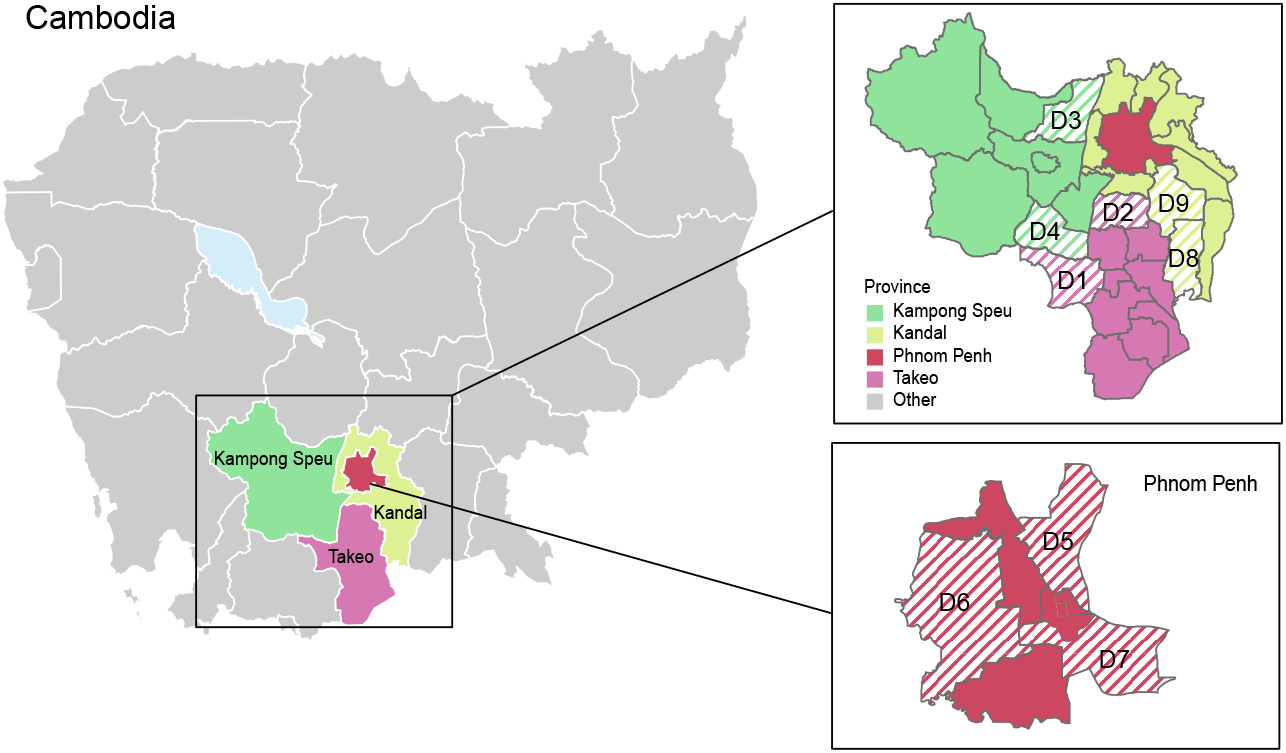
Geographical area of swine influenza surveillance study conducted in Cambodia, 2020–2022. Striped lines indicate the location by district (D1–D9) of sampled pig slaughterhouses in Kampong Speu, Kandal, Phnom Penh and Takeo provinces.

**Table 1.** Quantitative reverse transcription PCR (RT-qPCR) positivity rate and seroprevalence of influenza A viruses from pigs in Cambodia, 2020–2022.

| Sampling locations | RT-qPCR |  | IAV seroprevalence |  |
| --- | --- | --- | --- | --- |
|  | No. of nasal swab tested | No. of positive nasal swab (%) | No. of sera tested | No. of positive sera (%) |
| **Takeo Province** |  |  |  |  |
| District 1 | 769 | 17 (2.2%) | 760 | 201 (26.4%) |
| District 2 | 242 | 0 (0%) | 242 | 127 (52.5%) |
| sub-total | 1011 | 17 (1.7%) | 1002 | 328 (32.7%) |
| **Kampong Speu Province** |  |  |  |  |
| District 3 | 870 | 3 (0.3%) | 870 | 270 (31.0%) |
| District 4 | 382 | 0 (0%) | 379 | 149 (39.3%) |
| sub-total | 1252 | 3 (0.2%) | 1249 | 419 (33.5%) |
| **Phnom Penh Province** |  |  |  |  |
| District 5 | 539 | 2 (0.4%) | 538 | 56 (10.4%) |
| District 6 | 157 | 9 (5.7%) | 157 | 70 (44.6%) |
| District 7 | 391 | 8 (2.1%) | 391 | 178 (45.5%) |
| sub-total | 1087 | 19 (1.8%) | 1086 | 304 (30.0%) |
| **Kandal Province** |  |  |  |  |
| District 8 | 258 | 19 (7.4%) | 255 | 101 (39.6%) |
| District 9 | 481 | 14 (2.9%) | 477 | 169 (35.4%) |
| sub-total | 739 | 33 (4.5%) | 732 | 270 (36.9%) |
| **Total** | **4089** | **72 (1.78%)** | **4069** | **1321 (32.5%)** |

We obtained and analyzed complete or partial swine influenza genomes from 45 nasal swab samples (Table S1). Both H1 and H3 HA subtypes as well as N1 and N2 NA subtypes were identified during the sampling period, in combination with 10 HA and 9 NA individual co-infections (Fig. 3a). H1N1 subtype was predominant, being present in 37 (82.2%) of 45 sequenced pig samples (Fig. 3a). To trace the evolutionary origin of individual gene segments of swIAV from Cambodia, maximum likelihood phylogenies were reconstructed for each gene segment: H1-HA (n= 1,009), N1-NA (n= 986), H3-HA (n= 766), N2-NA (n= 773), PB2 (n= 924), PB1 (n= 923), PA (n= 915), NP (n= 927), MP (n= 927) and NS (n= 927).

**Fig. 3.**
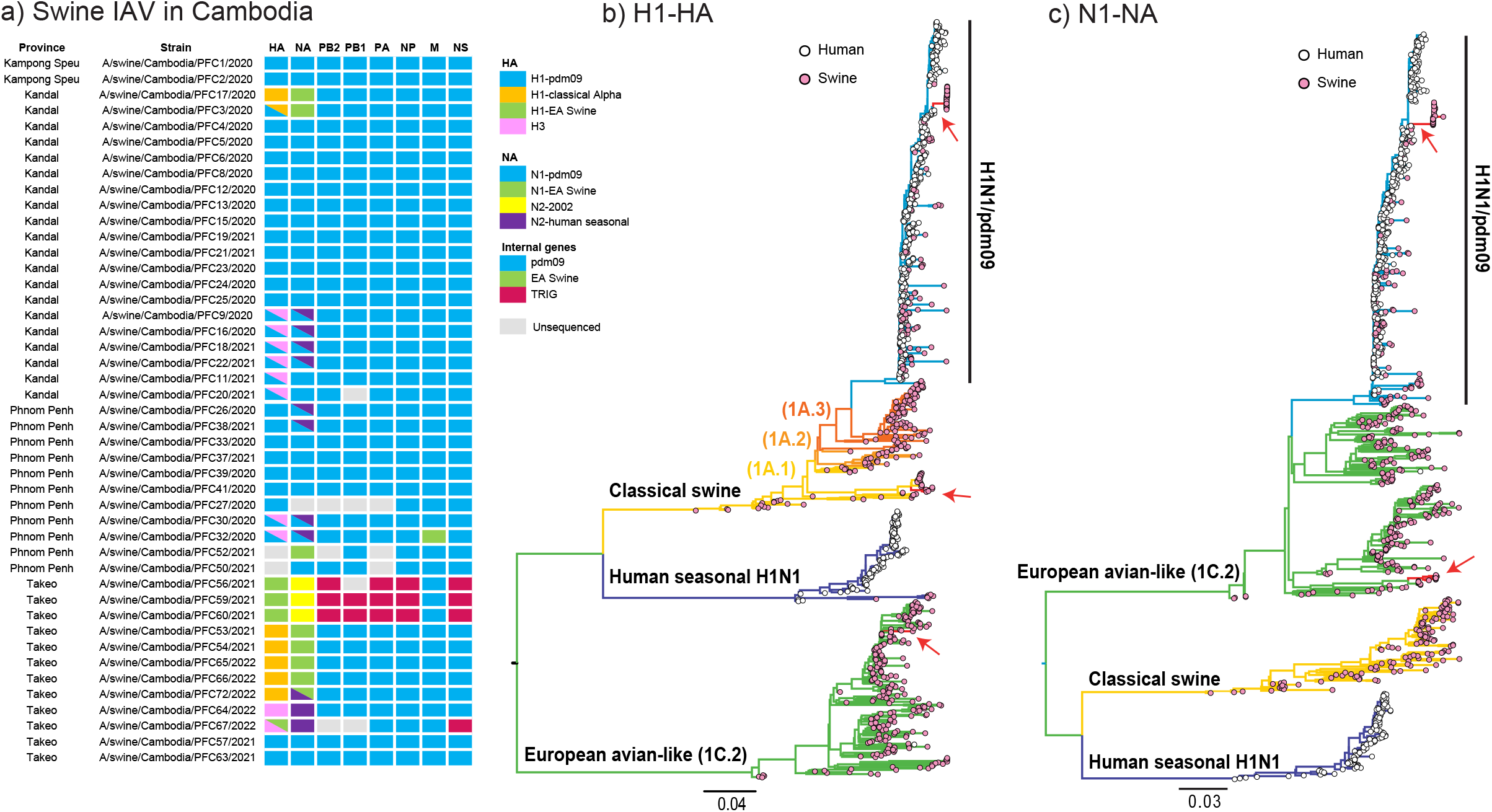
Diversity of swIAV strains. (a) Genomic constellations for each gene segment of swIAV strains from sampled pigs in Cambodia. Coloured boxes represent the major swIAV lineages detected in this study. (b, c) Evolutionary relationships of the H1-HA (b) and N1-NA (c) genes of human and swine influenza viruses inferred by maximum likelihood method in RAxML. Human and swine influenza viruses are denoted by pink and white tip circles, respectively. Coloured branches represent different IAV lineages, red branches and arrows indicate new swIAV gene sequences from this study.

Phylogenetic inference indicated that diverse lineages of H1 were in circulation within Cambodia. The majority of the new H1 and N1 sequences were derivatives of the human H1N1/pdm09 lineage (Fig. 3b and 3c). Genomic analysis revealed a unique reassortant European avian-like (EA) swine H1N2 subtype in four samples. This H1N2 subtype possessed an EA H1-HA gene (shown by green boxes/branches in Fig. 3a and 3b, Fig. 4a), while the acquisition of the N2 gene was from swine-like N2-2002 (denoted by yellow boxes in Fig. 3a) but nested within human seasonal N2. Classical swine (CS) H1N1 viruses were detected in seven swine samples, all with HA genes belonging to the CS H1 alpha lineage (orange boxes in Fig. 3a). Notably, these CS H1N1 viruses possessed the EA swine N1 gene (green boxes/branches in Fig. 3a and Fig. 3b). In one sample (A/swine/Cambodia/PFC3/2020) the HA of CS H1 and H1N1/pdm09 were both detected.

**Fig. 4.**
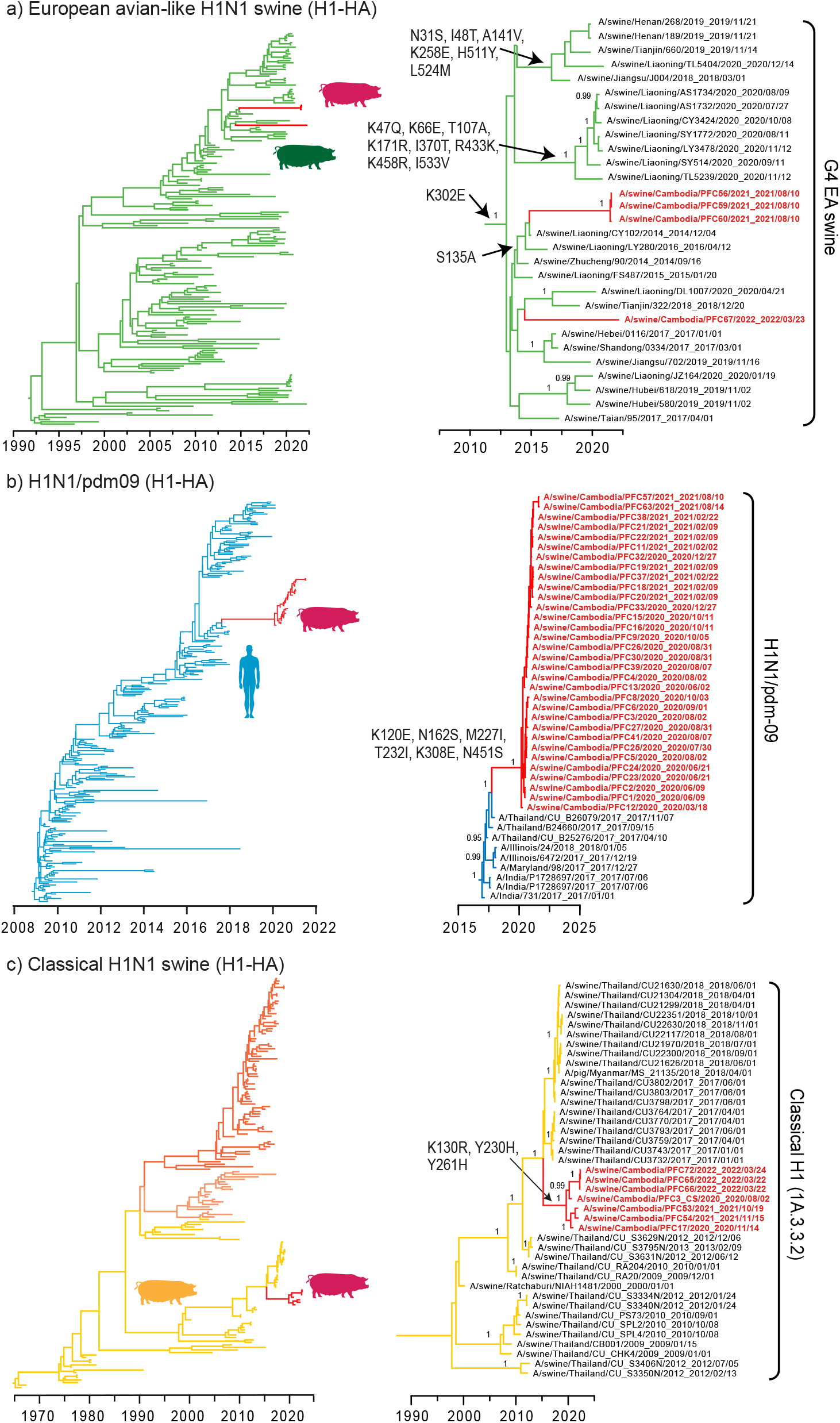
Time scaled phylogenies of H1-HA genes of human and swine influenza viruses. (a) European avian-like H1N1 swine viruses, represented by green branches. (b) human H1N1/pdm09 viruses, represented by blue branches. (c) Classical H1N1 swine viruses, represented by orange/yellow branches. Red branches represent swIAV sequences from this study. Persistent amino acid substitutions are indicated at the nodes. Bayesian posterior probability values (≥ 0.95) are shown at major nodes.

We detected H3 influenza subtypes in 10 (22.2%) of 45 sequenced pig samples from Cambodia. Eight swine samples were co-detected with H1N1/pdm09 and H3-HA viruses (shown by blue–pink gradient boxes in Fig. 3a). Six of these samples contained both an N2-NA gene and the N1 gene of H1N1/pdm09 virus (Fig. 3a). Another two samples (A/swine/Cambodia/PFC26/2020 and A/swine/Cambodia/PFC38/2021) showed evidence of H1N1/pdm09 and co-infection with H3N2 NA genes, but an H3 sequence was not obtained. These H1N1/pdm09 viruses were present in slaughterhouse pigs in all four provinces and H3N2 viruses were present in three provinces, in contrast to CS H1N1 and EA H1N2 subtypes which were mostly observed in Kandal and Takeo. As such, we found nine distinct swIAV HA and NA lineages are currently co-circulating in pig populations within Cambodia, facilitating the mixing of diverse genetic lineages through gene reassortment.

### Genesis of novel EA H1N2 reassortant virus in swine

We next unraveled the parental origin and gene constellations of the novel European avian-like H1N2 reassortant viruses. Four swine samples collected from Takeo Province (A/swine/Cambodia/PFC56/2021, A/swine/Cambodia/PFC59/2021, A/swine/Cambodia/PFC60/2021, A/swine/Cambodia/PFC67/2022) were assigned to EA H1N2 subtype (Fig. 3a and Fig. 4a). The N2 of EA H1N2 was most closely related to the North American swine N2-2002 lineage. In contrast, the internal genes of EA H1N2 subtype were acquired from the North American triple-reassortant internal gene (TRIG) cassette, except the MP gene that was derived from the H1N1/pdm09 lineage (Fig. 3a, *SI Appendix* Fig. S1–S6). The gene constellations are distinct from previously described EA swine viruses from Europe and China, that contain mostly H1N1/pdm09-origin internal genes, indicating that the contemporary EA H1N2 reasssortant viruses in Cambodia arose from diverse ancestral origins.

Our temporal phylogeny of the H1-HA gene showed that three out of four EA H1 swine sequences from Cambodia clustered to form a monophyletic group (posterior probability (PP)=1.00, denoted by red branches in Fig. 4a), with a mean TMRCA estimated around June 2021 (95% HPD intervals: April 2021–August 2021, Table 2). These three EA H1 sequences were most closely related to the EA H1N1 G4 genotype viruses found in swine from Liaoning, China between 2014–2016. The introduction of G4 H1-HA genes into pigs from Cambodia may have occurred between October 2014 and June 2021, up to 7 years before its first detection in Cambodia. Comparatively, one EA swine sample (A/swine/Cambodia/PFC67/2022) collected in 2022 appeared to be phylogenetically segregated and it was more related to recent viruses from Tianjin and Liaoning in 2018–2020, reflecting a possible independent introduction (Fig. 4a, *SI Appendix* Fig. S7).

**Table 2.** Estimated times to most recent common ancestor (TMRCA) of various swIAV lineages in pigs from Cambodia.

| Lineage | Time to most recent common ancestor (Cambodian lineage) |  |  | Time to most recent common ancestor (stem) |  |  | Undetected<br>years |
| --- | --- | --- | --- | --- | --- | --- | --- |
|  | Mean | 95% lower HPD | 95% upper HPD | Mean | 95% lower HPD | 95% upper HPD |  |
| H1 CS | 27 June 2019 | 06 November 2018 | 18 January 2020 | 21 January 2015 | 06 March 2014 | 09 December 2015 | 4.43 |
| H1 EA | 24 June 2021 | 01 April 2021 | 29 August 2021 | 04 October 2014 | 20 February 2014 | 02 December 2014 | 6.72 |
| H1 pdm09 | 16 February 2020 | 16 December 2019 | 16 March 2020 | 21 August 2017 | 22 May 2017 | 26 October 2017 | 2.49 |
| H3 seasonal group 1 | 09 February 2021 | 19 April 2019 | 10 November 2021 | 17 November 2010 | 20 December 2009 | 24 June 2011 | 10.23 |
| H3 seasonal group 2 | 01 July 2017 | 17 March 2015 | 23 April 2019 | 25 January 2012 | 25 March 2010 | 26 April 2013 | 5.43 |
| H3 seasonal group 3 | 14 March 2015 | 18 October 2012 | 02 December 2017 | 07 May 2011 | 26 April 2010 | 25 January 2012 | 3.85 |
| N1 EA swine | 06 November 2018 | 09 December 2017 | 03 August 2019 | 12 December 2014 | 08 October 2013 | 05 December 2015 | 3.90 |
| N1 pdm09 | 25 January 2020 | 30 October 2019 | 16 March 2020 | 21 November 2017 | 30 July 2017 | 27 February 2018 | 2.18 |
| N2 seasonal (EA<br>H1)* | 24 June 2021 | 25 March 2021 | 07 August 2021 | 28 November 2005 | 21 May 2004 | 26 May 2007 | 15.57 |
| N2 seasonal (CS H1)* | 09 August 2012 | 13 June 2010 | 10 August 2014 | 10 January 2008 | 17 March 2006 | 08 July 2009 | 4.58 |
| N2 seasonal (human<br>H3)* | 16 December 2021 | 03 August 2021 | 06 March 2022 | 06 May 2008 | 07 August 2006 | 06 November 2009 | 13.61 |

\*The corresponding HA of the N2 seasonal genes are indicated in Fig. 5e. Abbreviations: CS, classical swine; EA, European avian-like swine; pdm09, human H1N1/2009 pandemic.

The EA H1-HA viruses have acquired mutations at a mean rate of 3.77×10^−3^ substitutions per site per year (s/s/y), which was similar to the rates estimated for the H1 of H1N1/pdm09 and CS viruses (mean rates of 3.90×10^−3^ s/s/y and 3.37×10^−3^ s/s/y, respectively). Notably, the G4 EA H1-HA lineage has acquired sequential HA amino acid mutations since 2014. The HA mutation S135A defines the 2021–2022 Cambodian and related Liaoning EA viruses that circulated in 2014–2016 (Fig. 4a). Liaoning EA viruses from 2020 displayed extensive HA mutations (K47Q, K66E, T107A, K171R, I370T, R433K, K458R and I533V) than earlier strains (Fig. 4a), indicative of EA lineage diversification in Asia.

The N2 genes of Cambodian EA H1N2 viruses were ultimately derived from an established swine lineage and a distinct human seasonal N2 lineage (PP=1.00, Fig. 5e). Of note, the mean TMRCA was estimated around June 2021 (95% HPD intervals: March 2021– August 2021, Table 2), which overlaps with the TMRCA of EA H1 gene lineage in pigs from Cambodia. These N1 sequences are most closely related to ancestral swine H3N2 viruses circulated in 2012–2016 from Canada (e.g. A/swine/Manitoba/D0208/2013) and North America (e.g. A/swine/Iowa/13E045/2013), but with a combined TMRCA of 2005 (Fig. 5e). Our data suggests that the North American swine N2 segment may have been introduced into Asia over 15 years ago, highlighting decades of unsampled diversity due to a lack of systematic surveillance of swine influenza viruses in Southeast Asia.

**Fig. 5.**
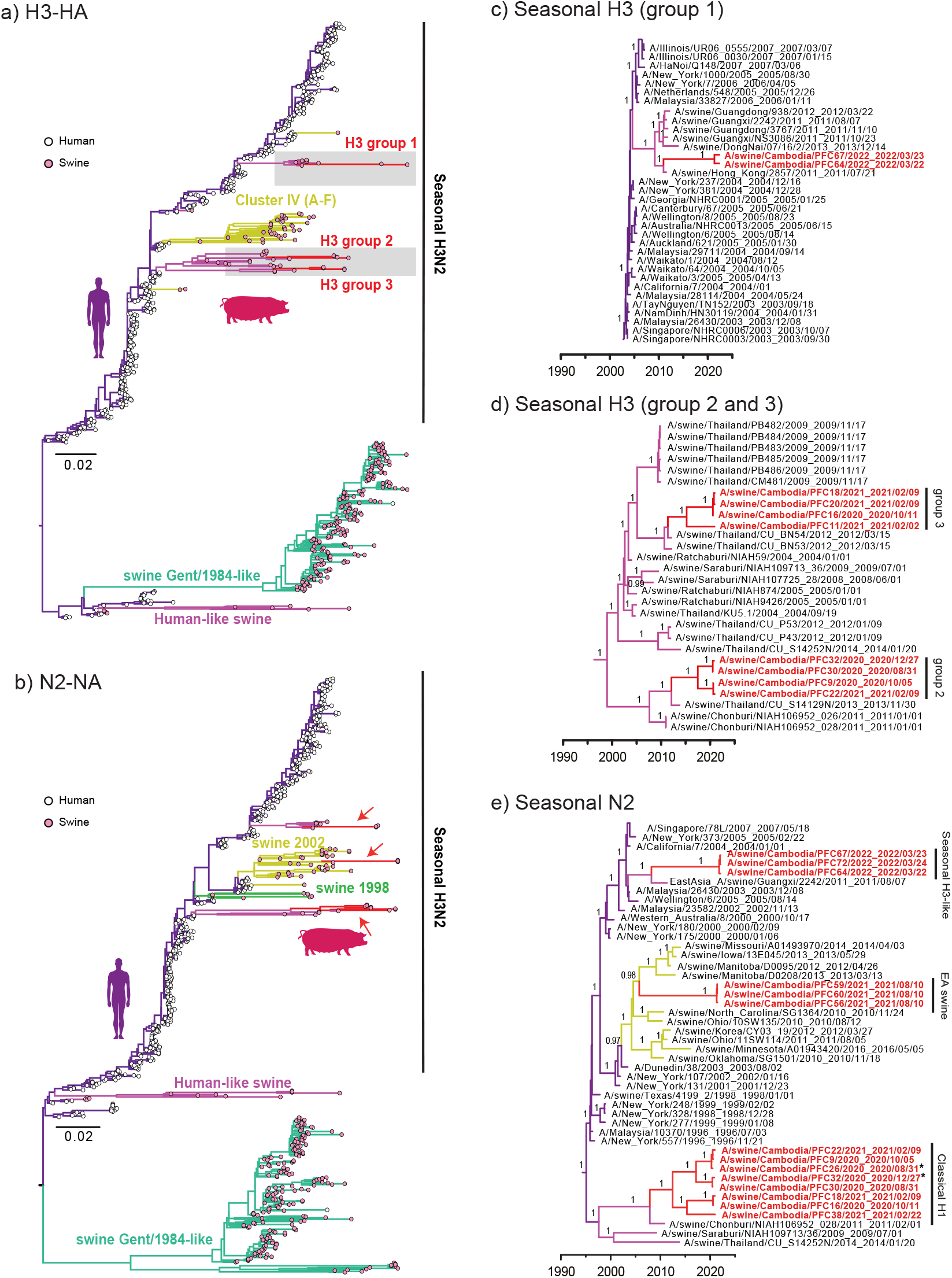
Phylogenies of H3-HA and N2-NA gene segments. (a, b) Evolutionary relationships of the H3-HA (a) and N2-NA (b) genes of influenza viruses inferred by maximum likelihood method in RAxML. Human and swine influenza viruses are denoted by pink and white tip circles, respectively. Coloured branches represent different IAV lineages. (c, d) Time scaled phylogenies of H3-HA genes. (e) Seasonal H3 avian-like H1N1 swine viruses. Red branches and arrows represent new swIAV sequences from this study. Bayesian posterior probability values (≥0.95) are shown at major nodes. *The corresponding HA gene of A/swine/Cambodia/PFC26/2020 is H1N1/pdm-09, whereas that of A/swine/Cambodia/PFC32/2020 contained both H1N1/pdm-09 and H3.

Most internal genes (PB2, PB1, PA, NP, MP and NS) of EA H1N2 virus from Cambodia pigs were found to have descended from North American TRIG lineage (Fig. 6). Phylogenetic positions of these internal genes of Cambodian EA viruses were either basal or nested within the CS H1N1 viral lineage. Of note, the NS gene of Cambodian EA viruses were related to the genes of G4 viruses from pigs in China, including recent Liaoning strains (e.g. A/swine/Liaoning/HLD1795/2020); these viruses collectively clustered within the TRIG lineage. In contrast, the MP gene of Cambodian EA viruses were most related to swine H1N1 viruses from China and North America, and nested within the human H1N1/pdm09 lineage. Interestingly, the H1, N1, most TRIG internal genes and MP gene of the three Cambodia EA swine exhibited overlapping TMRCA dates, estimated around October 2020 to August 2021 (*SI Appendix* Fig. S8). However, the associated PB2 showed an earlier TMRCA date in late 2017, indicative of independent introduction of the gene segment.

**Fig. 6.**
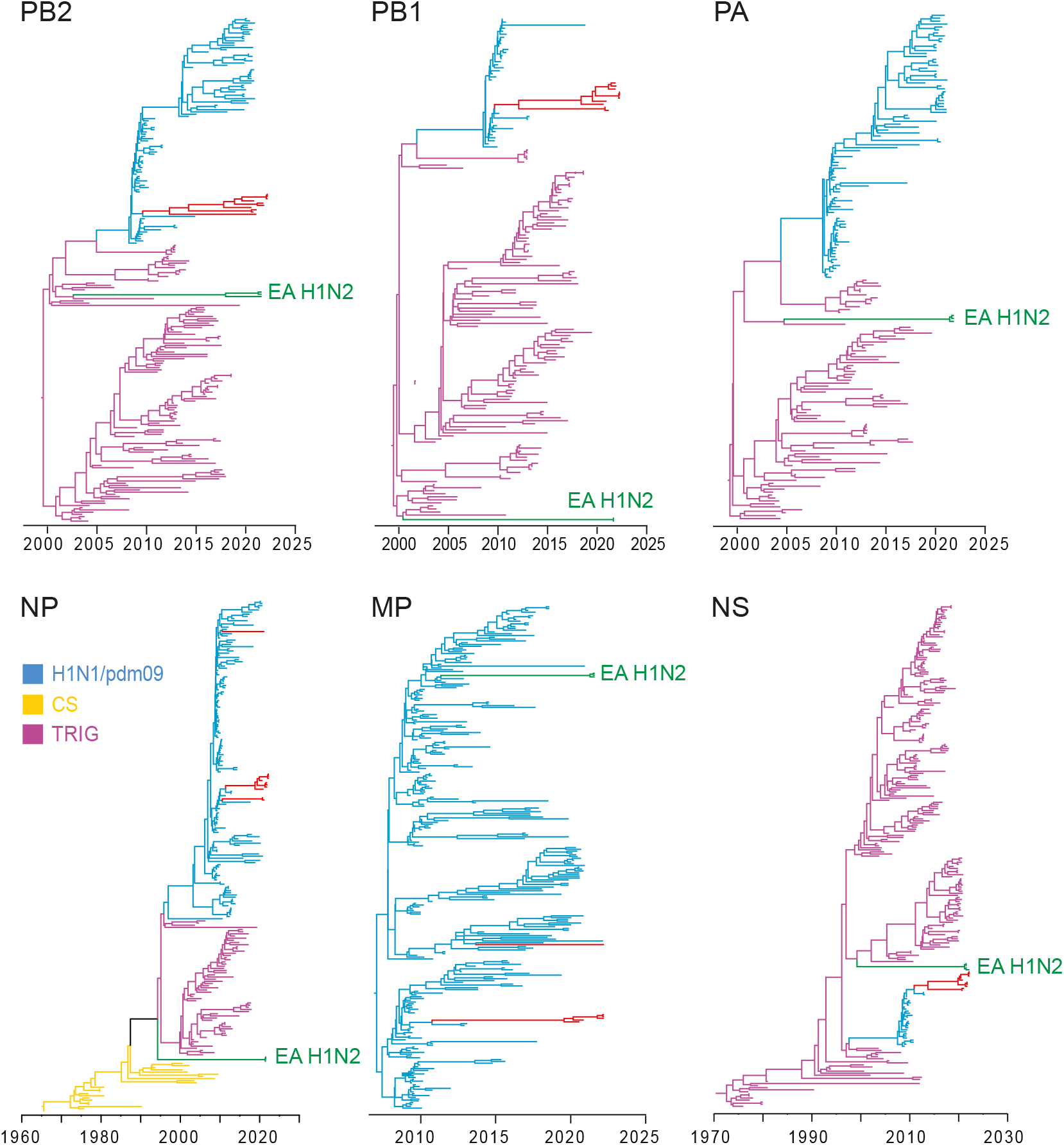
Temporal phylogenetic trees highlighting the origin of EA H1N2 internal genes. Purple, blue and orange branches represent TRIG, H1N1/pdm09 lineage and CS virus lineages, respectively. Red and green branches represent new swIAV and new EA H1N2 sequences from this study, respectively.

### Reverse zoonosis of H1N1/pdm09-like virus in pigs from Cambodia

Most of the H1 and N1 sequences from pigs in Cambodia belonged to human H1N1/pdm09 lineage, forming a distinct monophyletic sublineage (PP=1.00, H1 and N1 in Fig. 4b and *SI Appendix* Fig. S9a, respectively) that nested within the H1N1/pdm09 HA and NA lineages. The H1 and N1 gene segments had an estimated mean TMRCAs of approximately February 2020 and January 2020, respectively (both PP=1.00, denoted by red branches in Figs. 4b, *SI Appendix* Fig. S9a, Table 2). In the HA phylogeny, the most closely related strains are human H1N1/pdm09 viruses from Thailand collected in November 2017, and dating suggests that human-to-swine transmission occurred as early as August 2017, followed by enzootic spread of a genetically distinct H1N1/pdm09 sublineage among pigs in Cambodia. During this period, the swine H1N1/pdm09 viruses in Cambodia acquired six H1-HA amino acid substitutions (K120E, N162S, M227I, T232I, K308E and N451S) (Fig. 4b) and eight N1-NA substitutions (T9A, N50S, Q78K, P126H, I163V, T332I, I374V and V394I) (*SI Appendix* Fig. S9a). These HA and NA mutations may be due to adaptation to pigs following reverse zoonosis, however, it is not known if they may have arisen through virus circulation in Cambodian pigs or spread from other countries. The remaining internal genes (PB2, PB1, PA, NP, MP and NS) were predominantly derived from human H1N1/pdm09 viruses (*SI Appendix* Figs. S10–S15), except for A/swine/Cambodia/PFC32/2020 where the MP gene was from an EA H1N1 virus.

### Co-circulation of multiple classical swine H1N1 and H3N2 lineages in pig populations

Two enzootic swIAV viruses, CS H1 and H3N2, were also detected in pigs sampled from Kandal and Takeo provinces. The CS H1 subtype was detected in 7 swine samples, all HA sequences grouped into a monophyletic clade, which nested within the classical swine H1 alpha 1A.1.2 lineage (PP=1.00, Fig. 4c). The accompanying N1 gene is derived from EA viruses and the internal genes are from H1N1/pdm09 virus. The mean TMRCA of the Cambodian CS H1 sequences was estimated around June 2019 (Fig. 4c, Table 2) and they grouped with CS H1 viruses from Thailand detected from 2009–2018. The introduction of the CS H1 gene from Thailand to Cambodia likely occurred as early as January 2015. The EA N1 sequences of these samples (with an additional NA gene from A/swine/Cambodia/PFC52/2021) were most closely related to Thailand swine 2017 viruses and had a TMRCA of November 2018 (PP=1.00, *SI Appendix* Fig. S9b, Table 2), suggesting independent introduction into Cambodia than the slightly older CS H1 gene. These CS/EA H1N1 reassortants have also acquired amino acid mutations near the HA receptor binding sites (K130R, Y230H and Y261H) and in the NA protein (I53M, V62L and Q308H). The multiple origins of gene segments in these viruses highlights the frequent reassortment between diverse virus lineages that occurs in the region.

A high diversity of A/H3N2 subtype virus was found in 10 swine samples in Cambodia, with three distinct groups of H3N2 viruses circulating that all clustered with swine H3N2 viruses that were ultimately derived from human seasonal H3N2 (Fig. 5a and 5b). Interestingly, H3N2 virus was co-detected with H1N1/pdm09 HA genes in eight samples. Group 1 H3 contained two samples (A/swine/Cambodia/PFC64/2022 and A/swine/Cambodia/PFC67/2022), with the mean TMRCA estimated in February 2021 (Table 2). Both viruses were closely related to other swine H3 2011–2013 viruses from southern China (Guangdong and Guangxi) and southern Vietnam (Fig. 5c). Group 2 and 3 H3 each contained four viruses with estimated mean TMRCAs of July 2017 and March 2015, respectively (Fig. 5d, Table 2). Both groups were independently derived from Thailand swine viruses detected in swine since 2004. Prior to detection in swine, this H3N2 lineage was detected in humans as early as 1996, indicating there were multiple reverse-zoonoses of human H3N2 to swine that occurred between 1996 to 2004. The currently circulating swine N2 sequences from Cambodia also originated from human seasonal H3N2 viruses (Fig. 5e). Eight Cambodian swine N2 sequences formed a clade with an estimated TMRCA in August 2012 (Table 2) and are related to Thailand swIAVs from 2009–2014 (Fig. 5e). These N2 sequences may have been introduced from Thailand as early as January 2008.

### Spatiotemporal diffusion patterns of EA swine viruses

To determine the migration processes of EA swine H1-HA viruses, we reconstructed ancestral geographical locations and inferred diffusion patterns between locations using a discrete phylogeographic model. Phylogeographic reconstruction based on 1,073 sequences revealed that EA swine viruses circulated for decades among pigs in Europe until the early 2000s when we observed sporadic introductions into south central China and Southeast Asia (Fig. 7a, 7b). Since 2010, south central China became the dominant source for EA swine virus transmission, displaying strong diffusion links from south central China to east China, with strong migration support (BF>10,000, *SI Appendix* Table S2). Significant migration pathways with higher diffusion rates (2.11–3.83, Table S3) were also present from east China to other parts of China, including north China, northeast China, south central China (Fig. 7b). After 2016 there was an increased circulation of EA swine viruses between different regions of China, leading to a marked expansion of EA swine lineage diversity. During this period strong migration links were also evident from south central China to northeast China, and from northeast China to Southeast Asia (*SI Appendix* Table S2), including Cambodia. Nevertheless, phylogeographic interpretation should be treated with caution due to a lack of genetic data from Southeast Asia.

**Fig. 7.**
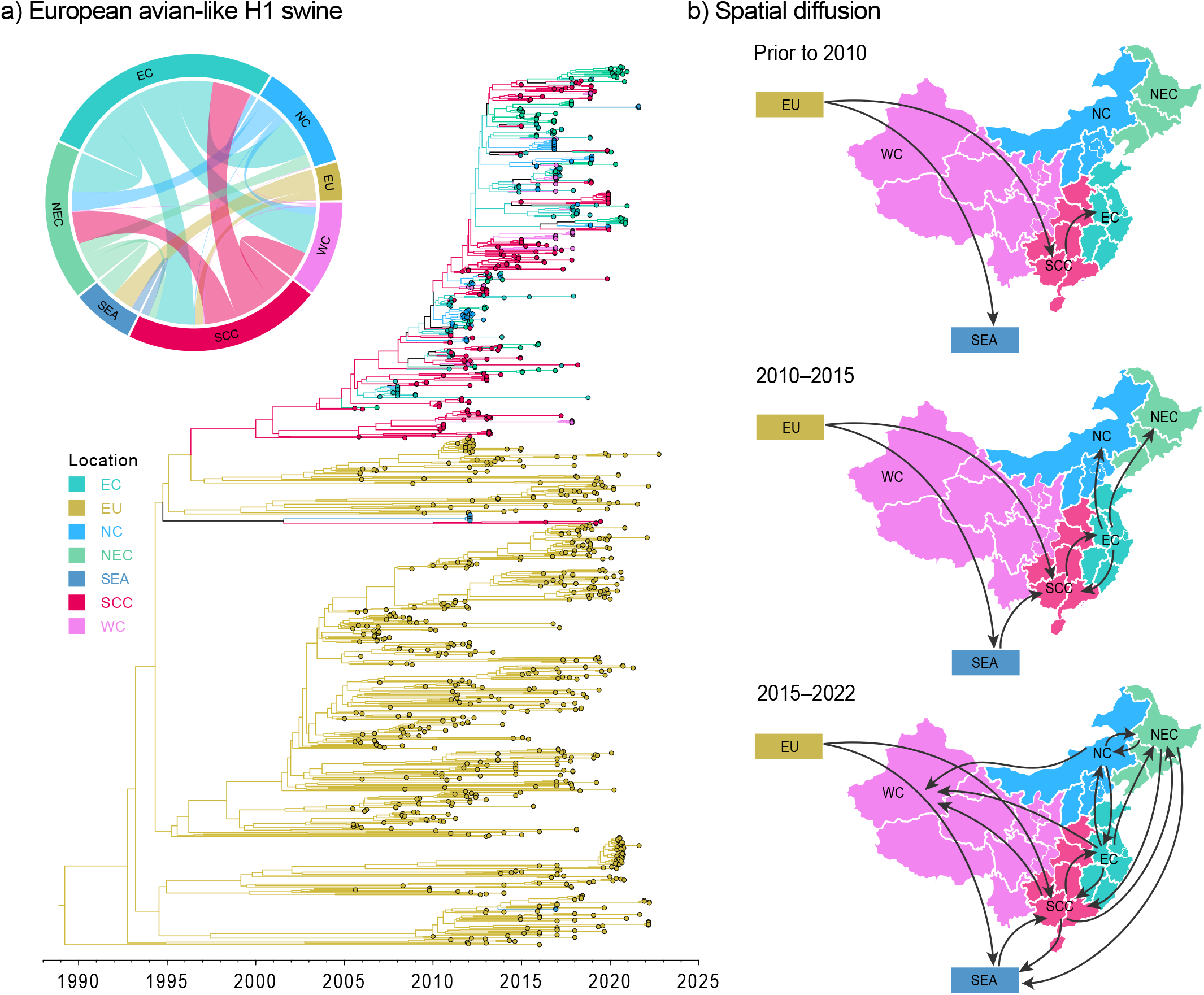
Discrete phylogeographic reconstruction of EA-like swine H1 diffusion dynamics. (a) Time-scaled maximum clade credibility tree of the Bayesian phylogeographic inference. Branch colour corresponds to geographic locations. The insert displays the circular migration plot based on log Bayes factor support between pairwise locations (see Supplementary Table S2 for support values). Abbreviations: EC, east China; EU, Europe; NC, north China; NEC, northeast China; SEA, Southeast Asia; SCC, south central China; WC, west China. (b) Spatial migration pathways of EA swine viruses through time.

## Discussion

Influenza A viruses are an integral part of a complex ecosystem that results in viral emergence and zoonotic diseases. Pigs are natural drivers for the emergence of novel influenza virus strains that cross species boundaries by means of shuffling gene segments between avian, swine and human hosts. While much is known about the genomic diversity of swIAVs in Europe and North America [32], the evolutionary dynamics of swIAV populations in Southeast Asia remains largely unknown.

Despite the difficulties posed by COVID-19, we conducted an extensive longitudinal study of influenza surveillance in pigs from slaughterhouses across four major provinces in Cambodia - spanning March 2020 through July 2022. Our study provides a detailed picture of virus circulation and dynamics in this region; showing diverse groups of swIAV viruses in pig populations and tracing the evolutionary footprints of their ancestral origins.

Our study highlights that circulating swIAV lineages in Southeast Asia acquired reassorted gene segments from diverse geographical origins, facilitated by inter- and intra-continental spread of previously segregated swIAV lineages, the same process that led to the emergence of the H1N1/pdm09 virus in Mexico [10]. This process also gave rise to the novel EA H1N2 subtype virus that likely emerged in late 2014, approximately 7 years before its first detection in Cambodia in 2021. This novel EA H1N2 has a unique gene constellation comprised of a swine-origin G4 EA H1-HA, a N2-HA originating from human viruses and internal genes from TRIG viruses. In contrast, the ancestral G4 EA H1N1 virus that emerged from pigs in Henan, China in 2011 [33], contained EA virus H1 and N1 genes, human H1N1/pdm09-derived internal genes and a TRIG-derived NS gene, and has since spread to other parts of China including Jilin, Liaoning and Tianjin [34]. The G4 EA virus HA binds preferentially to alpha 2,6-linked sialic acid receptors and replicates efficiently in human airway epithelial cells [33]. Given the zoonotic potential of the ancestral G4 EA H1N1 virus [34], the identification of this novel lineage in Cambodia is of particular importance, and highlights concerns surrounding inter-regional spread of swIAVs as a key driver in the emergence of new virus lineages. We also found multiple instances of CS H1 coupled with EA N1. Ancestral sequences to this viral lineage are primarily present in swine populations from Asia (China and Thailand) and North America, providing more evidence of global intermixing of previously distinct virus populations. Given the lack of pig surveillance, it is likely that this classical swine virus had circulated undetected in Cambodia since its introduction in 2017.

Pig value chains play a critical role in maintaining swine-to-swine virus transmission, regional and global migration, and the reassortment events that result from these activities. These prompt the establishment of diverse swIAV lineages among pig populations, including human influenza viruses that pose a constant threat for the reintroduction of antigenically and genetically distinct swIAVs being transmitted into humans [11, 32, 35, 36]. We found that H1N1/pdm09 virus is circulating enzootically in Cambodian swine herds. Since the H1N1/2009 pandemic, multiple H1N1/pdm09 virus spillovers back to pigs have occurred, seeding novel H1N1 reassortant viruses across the world [37-39]. We also detected multiple reverse zoonotic H3 lineages in Cambodian pigs that had circulated undetected for about 10 years. Follow-up experiments are needed to determine if the Cambodian H3 clades are antigenically distinct. The independent evolutionary trajectories of these viruses in pigs are recognized as a future pandemic threat [40]. Evidence of little to no antigenic cross-reactivity to currently circulating human influenza A viruses would indicate the potential for reintroduction of these viruses to susceptible human populations.

Pork is a major component of nutrition in many parts of Asia. The rapid growth in meat consumption is driven by increasing incomes, population sizes, and rapid urbanisation in developing countries [41]. However, a number of pig diseases such as influenza, porcine reproductive and respiratory syndrome (PRRS) and African swine fever (ASF) viruses have caused significant economic loss in swine industries [42]. As swine rearing operations in most countries continue to expand and intensify, routine and sustained surveillance in pigs is indispensable in identifying new viruses so that their zoonotic risk can be assessed. The current methods of disease surveillance are not fit for this purpose, primarily because sampling individual animals is expensive and time consuming. As most of the funding for this work in Southeast Asia comes from research grants, the efforts are also sporadic, which is reflected in the sparse swIAV genomic data available from the region. It is therefore critical that more efficient and continuous surveillance methods such as metagenomic surveillance [43] of air and wastewater samples in farms and slaughterhouses are integrated with automated analytical tools to rapidly provide information on changes in the spatio-temporal occurrence of a broad range of human and animal pathogens. Such a system would serve to improve animal health through selection of efficacious vaccines, and aid in human health by monitoring viruses with the potential for zoonotic transmission.

## Materials and Methods

### Ethics Statement

This study was approved by the ethics committees at LSHTM’s Institutional Review Board (approval number 16635) and Animal Welfare and Ethical Research Board (ref 2019-12), the National Ethics Committee for Health Research in Cambodia (ref 105), and the Human Research Protection Office (HRPO, ref A-21055) and Animal Care and Use Review Office (ACURO) of the USAMRDC Office of Research Protections. RNA samples were de-identified before transporting to the research laboratory at Duke-NUS.

### Sample collection, virus isolation and sequencing

From March 2020 to July 2022, we collected 4,089 nasal swab samples from pigs in 18 slaughterhouses located in four neighboring provinces (Kampong Speu, Kandal, Phnom Penh, and Takeo) of Cambodia (Fig. 2). Viral RNA was extracted from samples using the QIAmp Viral RNA Mini Kit (Qiagen) and real-time RT-PCR was performed to detect the presence of influenza A virus (M gene) using AgPath-ID™ One-Step RT-PCR Kit (Thermo Fisher Scientific). Positive influenza samples were then selected for whole genome sequencing using next generation sequencing (NGS) methods following previously published protocols [44]. Amplified cDNA was quantified using the Qubit dsDNA HS Assay kits (Thermo Fisher Scientific) and subsequently diluted to 1 ng/µL. Libraries were prepared using the Nextera XT DNA library preparation kits (Illumina) and pooled. The pooled libraries were sequenced using an illumina MiSeq sequencing platform at the Duke-NUS Genome Biology Facility, generating 250 bp paired-end reads. The NGS reads were checked with FastQC [45] in Unipro UGENE v40.1 [46] and were processed by trimming adaptors via Trimmomatic v0.39 [47]. For each sample, reads were de novo assembled using SPAdes v3.15.3 [48] and individual gene segment was determined by BLASTn v2.2.18 [49]. Reads were subsequently mapped to most similar segments using the map to reference tool in UGENE v40.1, and consensus sequences were extracted.

### Evolutionary analysis

All available global H1N1 IAV genomes (as of 12 January 2022) collected from human and swine hosts were downloaded from NCBI GenBank and the Global Initiative on Sharing All Influenza Data (GISAID) [50] databases during 1957–2022 (n > 15,000). The datasets were subsampled to 1,008 sequences for H1 and 766 sequences for H3 using SMOT [51]. Maximum likelihood (ML) phylogenies for H1, H3, N1, N2 and six internal genes (PB2, PB1, PA, NP, MP and NS) were individually reconstructed using RAxML-NG v1.1.0 [52] with the following number of sequences H1-HA (n= 1,009), N1-NA (n= 986), H3-HA (n= 766), N2-NA (n= 773), PB2 (n= 924), PB1 (n= 923), PA (n= 915), NP (n= 927), MP (n= 927) and NS (n= 927). Classification designations for H1 sequences were assigned using the Swine H1 Clade Classification tool provided by Influenza Research Database (IRD) [53]. H3 classifications were assigned based on phylogenetic inference [54, 55]. Lineages for internal genes were assigned using the OctoFLU pipeline [56] supplemented with A/swine/Arnsberg/1/1979 to indicate the EA-like lineage [57]. Percentage of nucleotide similarity within genetic lineages was measured using MEGA11 [58].

To estimate the date of introduction for each genomic segment into Cambodian pigs, we utilized the ML phylogenies reconstructed above to extract the Cambodian swine viruses and representative viruses that pertain to the corresponding lineages. Bayesian time-scaled trees were inferred for the HA and NA genes using BEAST v1.10.4 [59] with BEAGLE library v3.1.0 [60]. Posterior trees were sampled using an uncorrelated relaxed clock with lognormal distribution [61], under a generalized time-reversible (GTR) substitution model [62] with four rate categories of gamma-distributed site heterogeneity [63], and a Gaussian Markov random field (GMRF) tree prior with time-aware smoothing [64]. Two runs of Markov chain Monte Carlo sampling were performed, each with 100 million iterations and sampled every 10,000 trees. Convergence metrics were checked using Tracer v1.7.2 [65]. Time-scaled maximum clade credibility (MCC) trees were summarized after the removal of 10% burn-in using TreeAnnotator v1.10.4 [59]. The resulting MCC trees were visualized using FigTree v.1.4.2. Amino acid substitutions of the HA and NA were mapped using treesub (https://github.com/tamuri/treesub).

### Serological analysis

A total of 4,069 pig sera samples was collected in this study. The detection of influenza A nucleoprotein-specific antibody was assessed in duplicate using the ID Screen Influenza A Antibody Competition multi-species ELISA kits (IDVet-Innovative Diagnostic, France) according to the manufacturer’s protocol.

### Phylogeographic analysis

To examine the spatial patterns of EA-swine H1 viruses, we reconstructed the spatiotemporal pathways between locations using discrete phylogeographic analyses in BEAST v.10.4. All available H1 sequences of European avian-like swine viruses were downloaded from NCBI and GISAID databases (accessed 20 December 2022) and seven discrete geographical regions were coded (Europe, east China, north China, northeast China, south central China, west China and Southeast Asia). The HA dataset was subsampled using SMOF [51] and the final dataset consisted of 1,073 taxa. Asymmetric diffusion rates between discrete locations were inferred using Bayesian Stochastic Search Variable Selection (BSSVS) [66], strict molecular clock model, the HKY85 nucleotide substitution model with four rate categories of gamma-distributed site heterogeneity, and a coalescent exponential growth tree prior. At least four independent runs of 100 million generations were performed and sampled every 10,000 generations. The runs were combined after removal of burn-in, with effective sample size (ESS) of >200. Non-zero diffusion rates were calculated from the resulting log files. Bayes factor (BF) values were determined by SpreaD3 [67] and the circular migration diagram was plotted in R using the circlize package [68]. BF ≥1,000 indicates decisive support, 100≤BF<1000 as very strong support, 10≤BF<100 as strong support, and 3≤BF<100 as supported (*SI Appendix* Table S2).

## Acknowledgements

The study was supported by the United States (US) Department of Defense Threat Reduction Agency Cooperative Biological Research project: PigFluCam+ (HDTRA11810051), contracts HHSN272201400006C and 75N93021C00016 from the National Institute of Allergy and Infectious Diseases, National Institutes of Health, Department of Health and Human Services, USA, and by the Duke-NUS Signature Research Programme funded by the Ministry of Health, Singapore. We thank Dr Zhang Rong at Duke-NUS for providing technical support. We thank GISAID and all submitters for access to their databases and all submitters of data and sequences to these databases.

## Author contributions

J.W.R., G.J.D.S. and Y.C.F.S conceived the study; S.T., S.S., A.H. and H.H. designed field work and performed research; S.T., S.S., D.K., A.H., H.H., T.C. and C.M. collected samples and coordinated data; S.B. and C.S. conducted RT-PCR and ELISA; G.G.K.N and Z.Y. conducted sequencing; M.A.Z. designed and implemented pipelines for genomic processing; M.A.Z., J.M., F.Y.W. and Y.C.F.S analysed data; M.A.Z., J.W.R., G.J.D.S and Y.C.F.S. wrote the manuscript. All authors commented and reviewed the manuscript.

## Additional information

Accession codes: The new sequences generated in this study have been deposited in GenBank database under accession numbers XXXXXX to XXXXXX.

## References

1. Garten, R.J., et al., Antigenic and genetic characteristics of swine-origin 2009 A (H1N1) influenza viruses circulating in humans. Science, 2009. 325(5937): p. 197–201.

2. Smith, G.J., et al., Dating the emergence of pandemic influenza viruses. Proceedings of the National Academy of Sciences, 2009. 106(28): p. 11709–11712.

3. Vijaykrishna, D., et al., Long-term evolution and transmission dynamics of swine influenza A virus. Nature, 2011. 473(7348): p. 519–522.

4. VanderWaal, K. and J. Deen, Global trends in infectious diseases of swine. Proceedings of the National Academy of Sciences, 2018. 115(45): p. 11495–11500.

5. Davies, P.R., Intensive swine production and pork safety. Foodborne Pathogens and disease, 2011. 8(2): p. 189–201.

6. Fao, F., Food and agriculture organization of the United Nations. Rome, URL: http://faostat.fao.org, 2022.

7. Tep, B., Swine Production and Disease Status in Cambodia, in Third Regional Workshop on Swine Disease Control in Asia. 2018 General Directorate of Animal Health and Production Ministry of Agriculture, Forestry and Fisheries Cebu, Philippines

8. Smith, G.J., et al., Origins and evolutionary genomics of the 2009 swine-origin H1N1 influenza A epidemic. Nature, 2009. 459(7250): p. 1122–1125.

9. Fraser, C., et al., Pandemic potential of a strain of influenza A (H1N1): early findings. science, 2009. 324(5934): p. 1557–1561.

10. Mena, I., et al., Origins of the 2009 H1N1 influenza pandemic in swine in Mexico. eLife, 2016. 5: p. e16777.

11. Nelson, M.I., et al., Global migration of influenza A viruses in swine. Nature communications, 2015. 6.

12. Nasamran, C., et al., Persistence of pdm2009-H1N1 internal genes of swine influenza in pigs, Thailand. Scientific reports, 2020. 10(1): p. 1–12.

13. Charoenvisal, N., et al., Genetic characterization of Thai swine influenza viruses after the introduction of pandemic H1N1 2009. Virus genes, 2013. 47(1): p. 75–85.

14. Nonthabenjawan, N., et al., Genetic diversity of swine influenza viruses in Thai swine farms, 2011–2014. Virus Genes, 2015. 50(2): p. 221–230.

15. Takemae, N., et al., Antigenic variation of H1N1, H1N2 and H3N2 swine influenza viruses in Japan and Vietnam. Archives of virology, 2013. 158(4): p. 859–876.

16. Ngo, L.T., et al., Isolation of novel triple-reassortant swine H3N2 influenza viruses possessing the hemagglutinin and neuraminidase genes of a seasonal influenza virus in Vietnam in 2010. Influenza and Other Respiratory Viruses, 2012. 6(1): p. 6–10.

17. Baudon, E., et al., Detection of novel reassortant Influenza A (H3N2) and H1N1 2009 pandemic viruses in swine in Hanoi, Vietnam. Zoonoses and public health, 2015. 62(6): p. 429–434.

18. Mon, P.P., et al., Swine influenza viruses and pandemic H1N1-2009 infection in pigs, Myanmar. Transboundary and Emerging Diseases, 2020. 67(6): p. 2653–2666.

19. Long, J.S., et al., Host and viral determinants of influenza A virus species specificity. Nature Reviews Microbiology, 2019. 17(2): p. 67–81.

20. A/H5, W.C.o.t.W.H.O.C.o.H.I., Avian influenza A (H5N1) infection in humans. New England Journal of Medicine, 2005. 353(13): p. 1374–1385.

21. Chen, Y., et al., Human infections with the emerging avian influenza A H7N9 virus from wet market poultry: clinical analysis and characterisation of viral genome. The Lancet, 2013. 381(9881): p. 1916–1925.

22. Van Reeth, K., Avian and swine influenza viruses: our current understanding of the zoonotic risk. Veterinary research, 2007. 38(2): p. 243–260.

23. Myers, K.P., C.W. Olsen, and G.C. Gray, Cases of swine influenza in humans: a review of the literature. Clinical infectious diseases, 2007. 44(8): p. 1084–1088.

24. Zeller, M.A., et al., Complete Genome Sequences of Two Novel Human-Like H3N2 Influenza A Viruses, A/swine/Oklahoma/65980/2017 (H3N2) and A/Swine/Oklahoma/65260/2017 (H3N2), Detected in Swine in the United States. Microbiol Resour Announc, 2018. 7(20): p. e01203–18.

25. Rajao, D.S., et al., Novel Reassortant Human-Like H3N2 and H3N1 Influenza A Viruses Detected in Pigs Are Virulent and Antigenically Distinct from Swine Viruses Endemic to the United States. J Virol, 2015. 89(22): p. 11213–22.

26. Vincent, A.L., et al., Characterization of a newly emerged genetic cluster of H1N1 and H1N2 swine influenza virus in the United States. Virus Genes, 2009. 39(2): p. 176–85.

27. Zhou, N.N., et al., Genetic reassortment of avian, swine, and human influenza A viruses in American pigs. Journal of virology, 1999. 73(10): p. 8851–8856.

28. Webby, R.J., et al., Evolution of swine H3N2 influenza viruses in the United States. Journal of virology, 2000. 74(18): p. 8243–8251.

29. Chea, B., et al., Assessment of Pig Disease Prevention of Smallholder Farmers and Village Animal Health Workers in Rural and Peri-Urban Cambodia. Open Journal of Animal Sciences, 2020. 10(3): p. 572–591.

30. Scholtissek, C., Pigs as ‘mixing vessels’ for the creation of new pandemic influenza A viruses. Medical Principles and Practice, 1990. 2(2): p. 65–71.

31. Pensaert, M., et al., Evidence for the natural transmission of influenza A virus from wild ducks to swine and its potential importance for man. Bulletin of the World Health Organization, 1981. 59(1): p. 75.

32. Henritzi, D., et al., Surveillance of European Domestic Pig Populations Identifies an Emerging Reservoir of Potentially Zoonotic Swine Influenza A Viruses. Cell Host Microbe, 2020. 28(4): p. 614–627.e6.

33. Sun, H., et al., Prevalent Eurasian avian-like H1N1 swine influenza virus with 2009 pandemic viral genes facilitating human infection. Proceedings of the National Academy of Sciences, 2020. 117(29): p. 17204–17210.

34. Li, H., et al., Prevalence, Genetics and Evolutionary Properties of Eurasian Avian-like H1N1 Swine Influenza Viruses in Liaoning. Viruses, 2022. 14(3): p. 643.

35. Nelson, M.I., et al., Global transmission of influenza viruses from humans to swine. J Gen Virol, 2012. 93(Pt 10): p. 2195–2203.

36. Nelson, M.I., et al., Spatial dynamics of human-origin H1 influenza A virus in North American swine. PLoS Pathog, 2011. 7(6): p. e1002077.

37. Danilenko, D.M., et al., Antigenic and Genetic Characterization of Swine Influenza Viruses Identified in the European Region of Russia, 2014-2020. Front Microbiol, 2021. 12: p. 662028.

38. Ryt-Hansen, P., et al., Co-circulation of multiple influenza A reassortants in swine harboring genes from seasonal human and swine influenza viruses. Elife, 2021. 10.

39. Mon, P.P., et al., Swine influenza viruses and pandemic H1N1-2009 infection in pigs, Myanmar. Transbound Emerg Dis, 2020. 67(6): p. 2653–2666.

40. Wong, F.Y.K., et al., Divergent Human-Origin Influenza Viruses Detected in Australian Swine Populations. J Virol, 2018. 92(16).

41. Mottet, A. and G. Tempio, Global poultry production: current state and future outlook and challenges. World’s Poultry Science Journal, 2017. 73(2): p. 245–256.

42. Woonwong, Y., D. Do Tien, and R. Thanawongnuwech, The Future of the Pig Industry After the Introduction of African Swine Fever into Asia. Animal Frontiers, 2020. 10(4): p. 30–37.

43. Holmes, E.C., The Ecology of Viral Emergence. Annu Rev Virol, 2022. 9(1): p. 173–192.

44. Ali, M., et al., Genetic characterization of highly pathogenic avian influenza A (H5N8) virus in Pakistani live bird markets reveals rapid diversification of clade 2.3. 4.4 b viruses. Viruses, 2021. 13(8): p. 1633.

45. Andrews, S., FastQC: a quality control tool for high throughput sequence data. 2010.

46. Okonechnikov, K., O. Golosova, and M. Fursov, Unipro UGENE: a unified bioinformatics toolkit. Bioinformatics, 2012. 28(8): p. 1166–7.

47. Bolger, A.M., M. Lohse, and B. Usadel, Trimmomatic: a flexible trimmer for Illumina sequence data. Bioinformatics, 2014. 30(15): p. 2114–2120.

48. Bankevich, A., et al., SPAdes: a new genome assembly algorithm and its applications to single-cell sequencing. Journal of computational biology, 2012. 19(5): p. 455–477.

49. Altschul, S.F., et al., Gapped BLAST and PSI-BLAST: a new generation of protein database search programs. Nucleic acids research, 1997. 25(17): p. 3389–3402.

50. Shu, Y. and J. McCauley, GISAID: Global initiative on sharing all influenza data– from vision to reality. Eurosurveillance, 2017. 22(13): p. 30494.

51. Arendsee, Z.W., A.L. Vincent, and T.K. Anderson, smot: a python package and CLI tool for contextual.

52. Kozlov, A.M., et al., RAxML-NG: a fast, scalable and user-friendly tool for maximum likelihood phylogenetic inference. Bioinformatics, 2019. 35(21): p. 4453–4455.

53. Anderson, T.K., et al., A Phylogeny-Based Global Nomenclature System and Automated Annotation Tool for H1 Hemagglutinin Genes from Swine Influenza A Viruses. mSphere, 2016. 1(6): p. e00275–16.

54. Anderson, T.K., et al., Swine influenza A viruses and the tangled relationship with humans. Cold Spring Harbor Perspectives in Medicine, 2021. 11(3): p. a038737.

55. Watson, S.J., et al., Molecular epidemiology and evolution of influenza viruses circulating within European swine between 2009 and 2013. Journal of virology, 2015. 89(19): p. 9920–9931.

56. Chang, J., et al., octoFLU: Automated Classification for the Evolutionary Origin of Influenza A Virus Gene Sequences Detected in US Swine. Microbiology resource announcements, 2019. 8(32): p. e00673–19.

57. Zell, R., C. Scholtissek, and S. Ludwig, Genetics, evolution, and the zoonotic capacity of European Swine influenza viruses. Curr Top Microbiol Immunol, 2013. 370: p. 29–55.

58. Kumar, S., et al., MEGA X: molecular evolutionary genetics analysis across computing platforms. Molecular biology and evolution, 2018. 35(6): p. 1547–1549.

59. Drummond, A.J., et al., Bayesian phylogenetics with BEAUti and the BEAST 1.7. Molecular biology and evolution, 2012. 29(8): p. 1969–1973.

60. Ayres, D.L., et al., BEAGLE: an application programming interface and high-performance computing library for statistical phylogenetics. Systematic biology, 2011. 61(1): p. 170–173.

61. Drummond, A.J., et al., Relaxed phylogenetics and dating with confidence. PLoS biology, 2006. 4(5): p. e88.

62. Tavaré, S., Some probabilistic and statistical problems in the analysis of DNA sequences. Lectures on mathematics in the life sciences, 1986. 17(2): p. 57–86.

63. Yang, Z., Maximum likelihood phylogenetic estimation from DNA sequences with variable rates over sites: approximate methods. Journal of Molecular evolution, 1994. 39(3): p. 306–314.

64. Minin, V.N., E.W. Bloomquist, and M.A. Suchard, Smooth skyride through a rough skyline: Bayesian coalescent-based inference of population dynamics. Molecular biology and evolution, 2008. 25(7): p. 1459–1471.

65. Rambaut, A. and A.J. Drummond, Tracer v1. 6 http://beast.bio.ed.ac.uk. 2007, Tracer.

66. Lemey, P., et al., Bayesian phylogeography finds its roots. PLoS Comput Biol, 2009. 5(9): p. e1000520.

67. Bielejec, F., et al., SpreaD3: Interactive Visualization of Spatiotemporal History and Trait Evolutionary Processes. Mol Biol Evol, 2016. 33(8): p. 2167–9.

68. Gu, Z., et al., circlize Implements and enhances circular visualization in R. Bioinformatics, 2014. 30(19): p. 2811–2.

