## Supplementary material for "The genomic landscape of swine influenza A viruses in Southeast Asia"

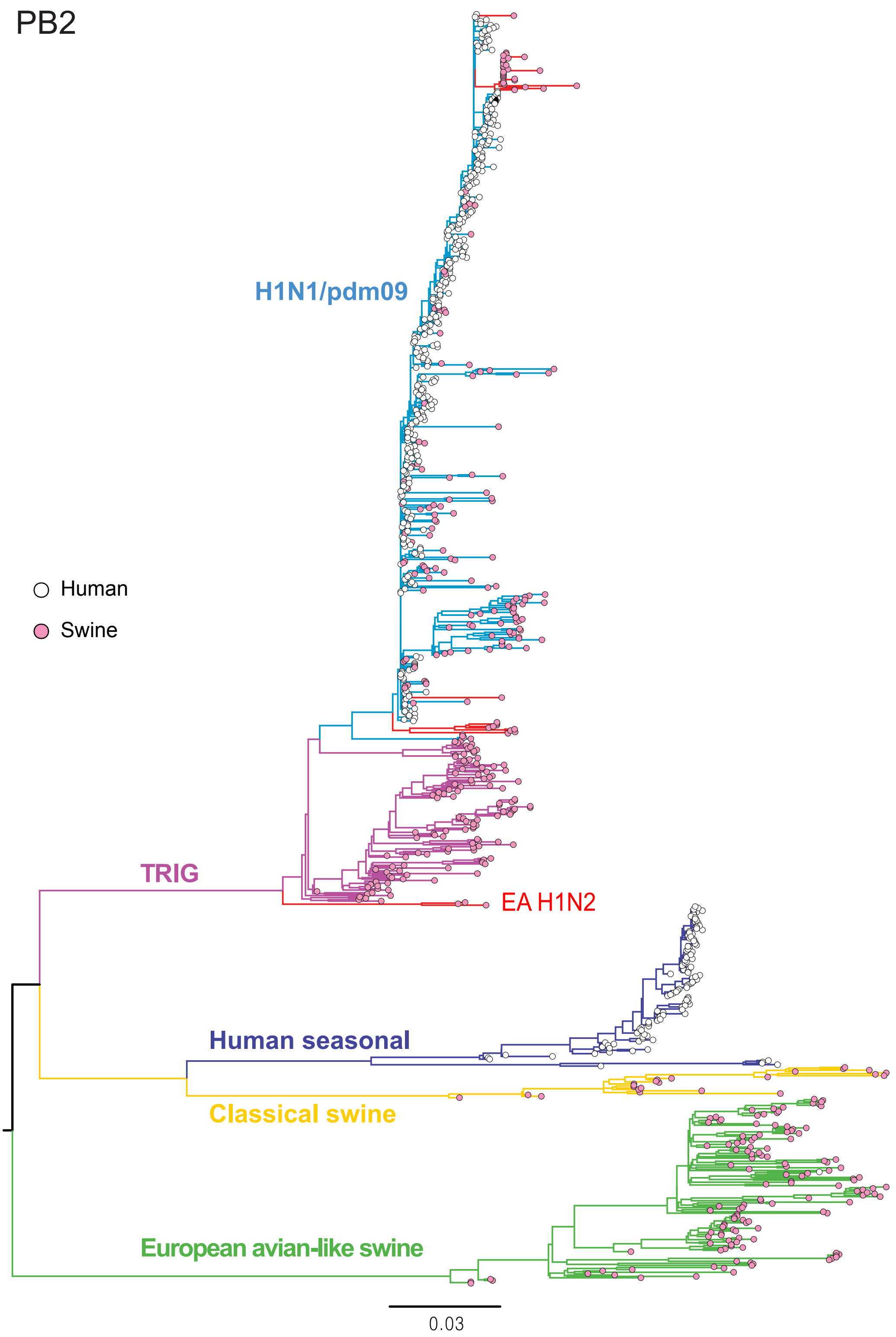


**Fig. S1**. Maximum likelihood phylogeny of the PB2 gene of human and swine influenza viruses. Coloured branches represent major lineages. Red branches denote swine sequences generated in this study. The novel EA H1N2 from pigs in Cambodia is indicated. Human and swine viruses are shown by pink and white tip circles. The scale bar represents the number of nucleotide substitutions per site.

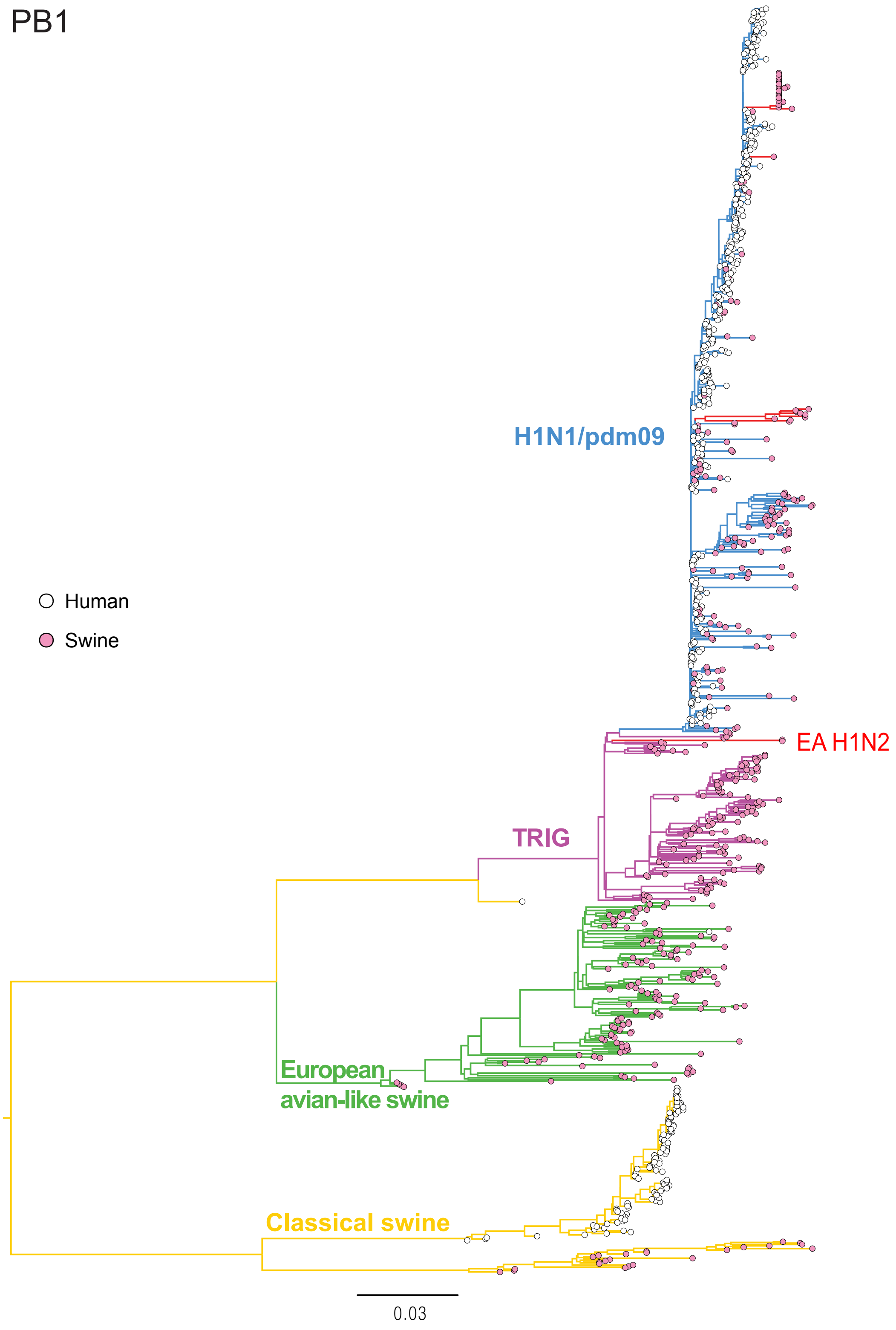


**Fig. S2**. Maximum likelihood phylogeny of the PB1 gene of human and swine influenza viruses. Coloured branches represent major lineages. Red branches denote swine sequences generated in this study. The novel EA H1N2 from pigs in Cambodia is indicated. Human and swine viruses are shown by pink and white tip circles. The scale bar represents the number of nucleotide substitutions per site.

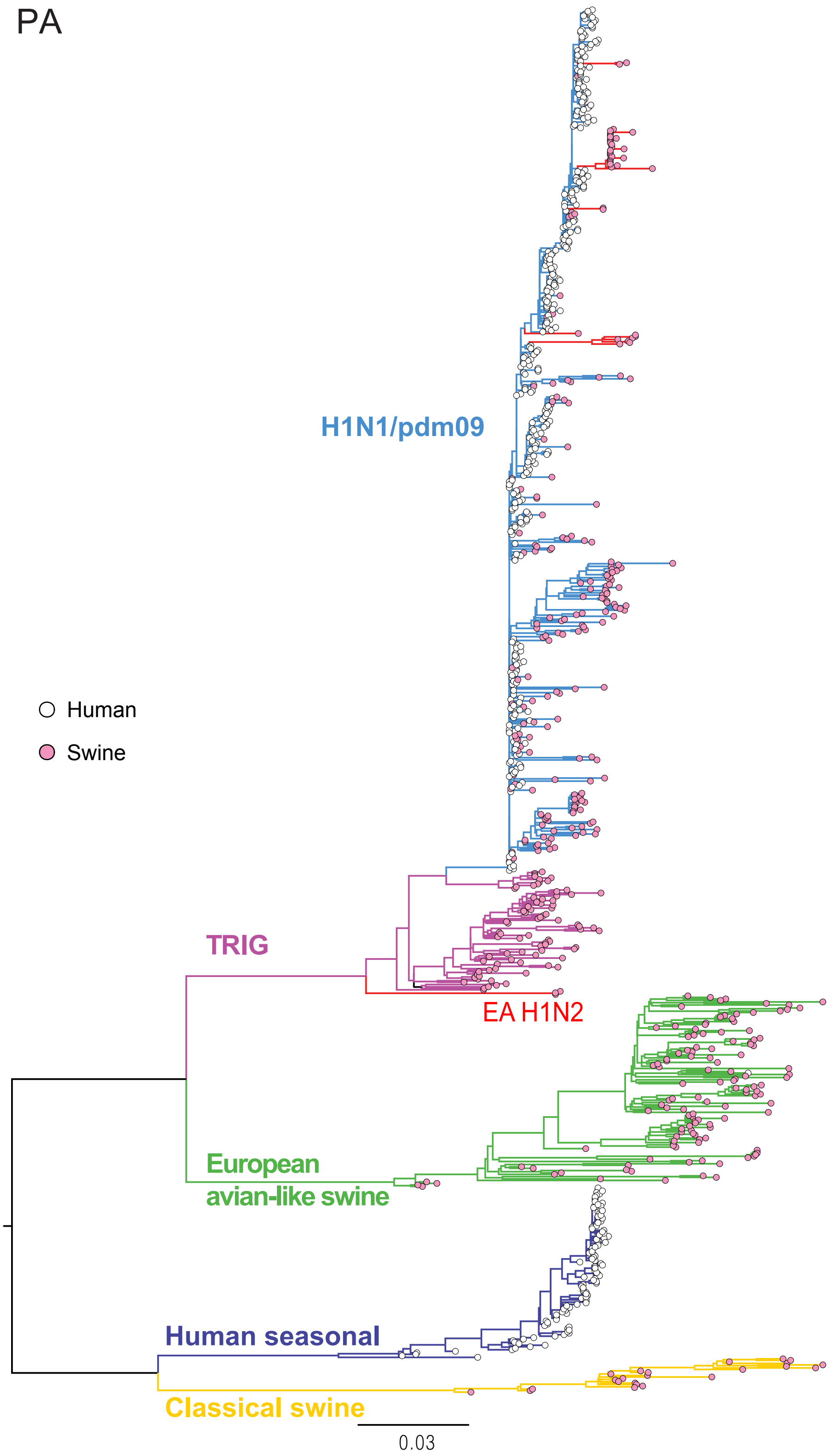


**Fig. S3**. Maximum likelihood phylogeny of the PA gene of human and swine influenza viruses. Coloured branches represent major lineages. Red branches denote swine sequences generated in this study. The novel EA H1N2 from pigs in Cambodia is indicated. Human and swine viruses are shown by pink and white tip circles. The scale bar represents the number of nucleotide substitutions per site.

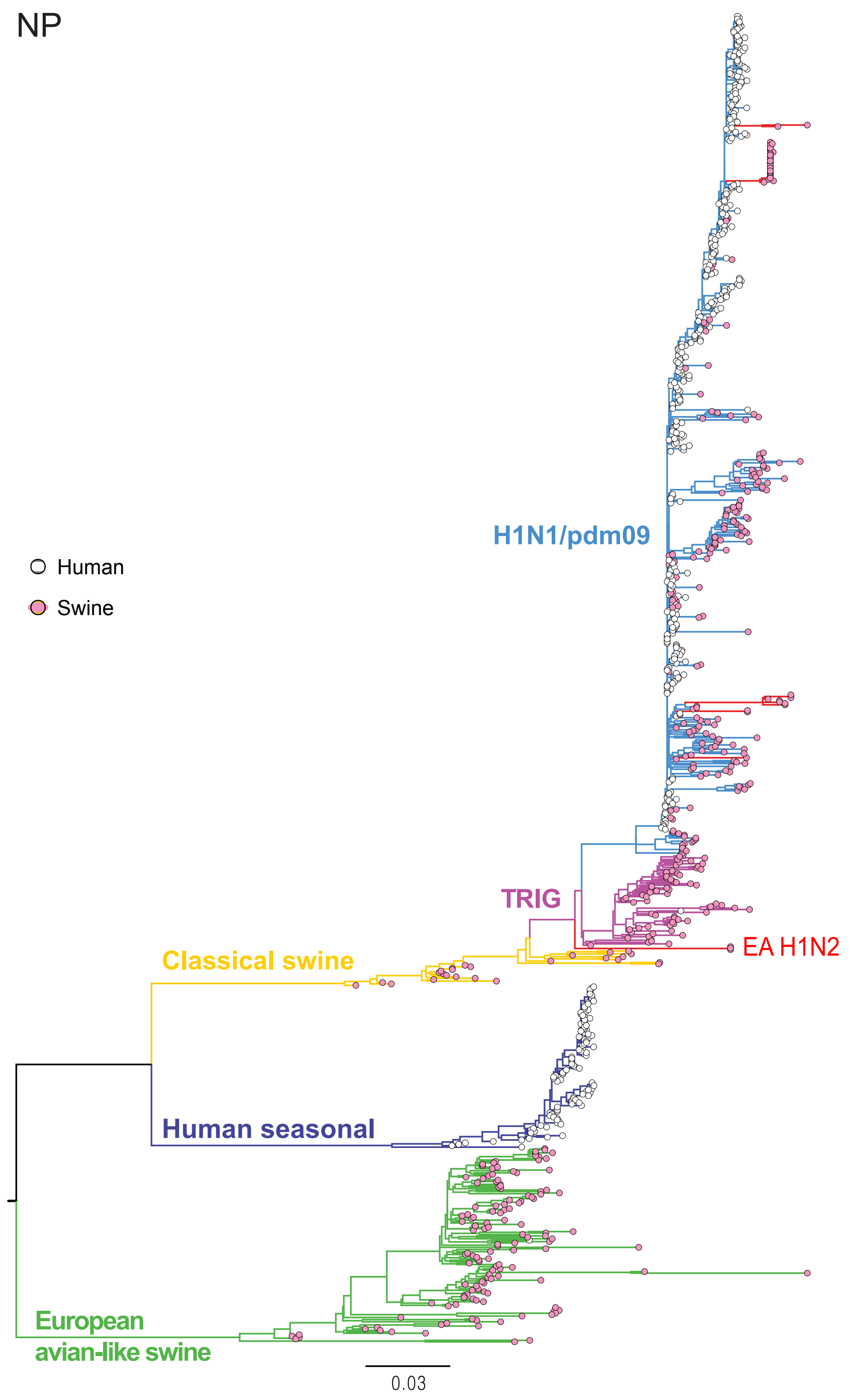


**Fig. S4**. Maximum likelihood phylogeny of the NP gene of human and swine influenza viruses. Coloured branches represent major lineages. Red branches denote swine sequences generated in this study. The novel EA H1N2 from pigs in Cambodia is indicated. Human and swine viruses are shown by pink and white tip circles. The scale bar represents the number of nucleotide substitutions per site.

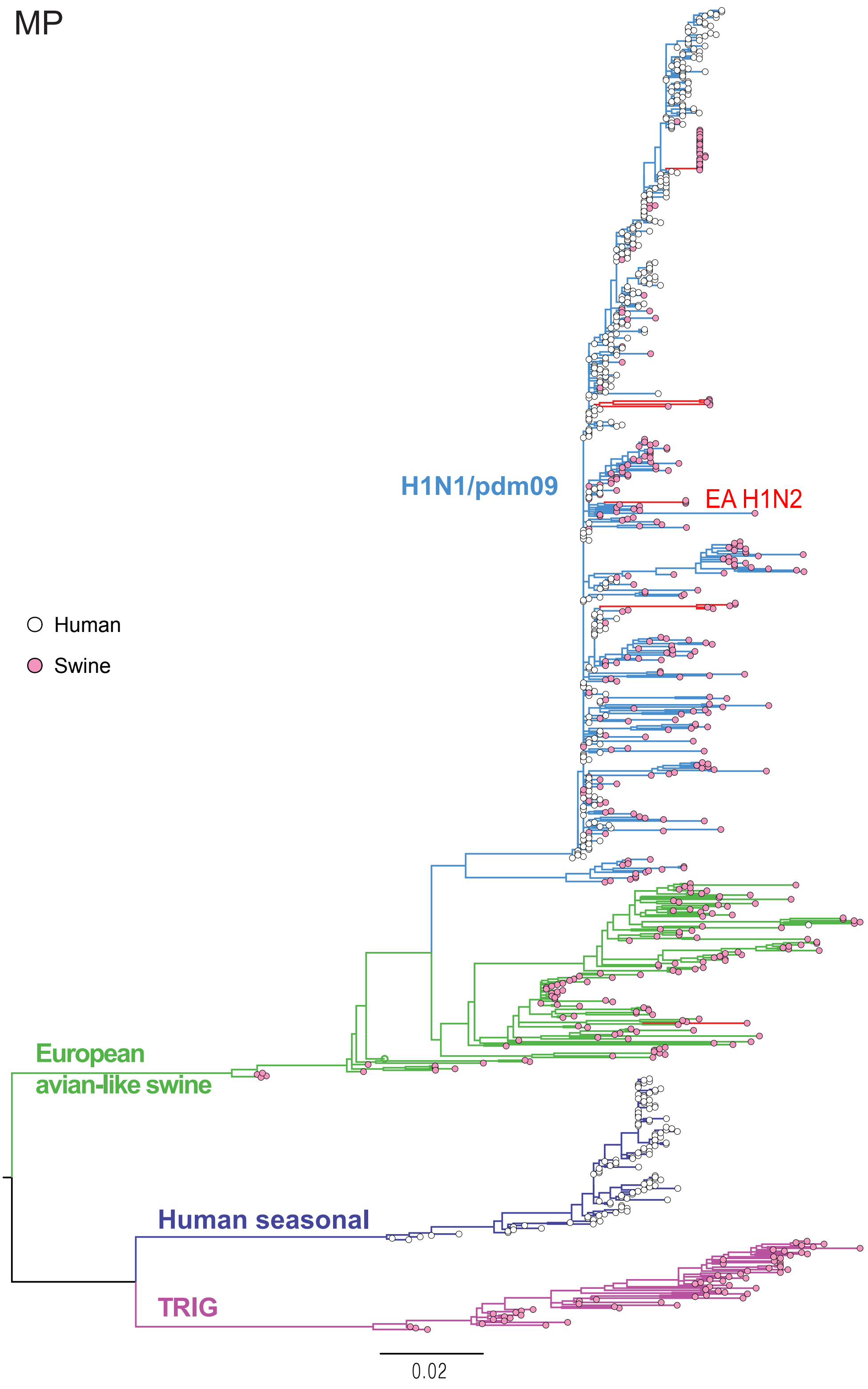


**Fig. S5**. Maximum likelihood phylogeny of the MP gene of human and swine influenza viruses. Coloured branches represent major lineages. Red branches denote swine sequences generated in this study. The novel EA H1N2 from pigs in Cambodia is indicated. Human and swine viruses are shown by pink and white tip circles. The scale bar represents the number of nucleotide substitutions per site.

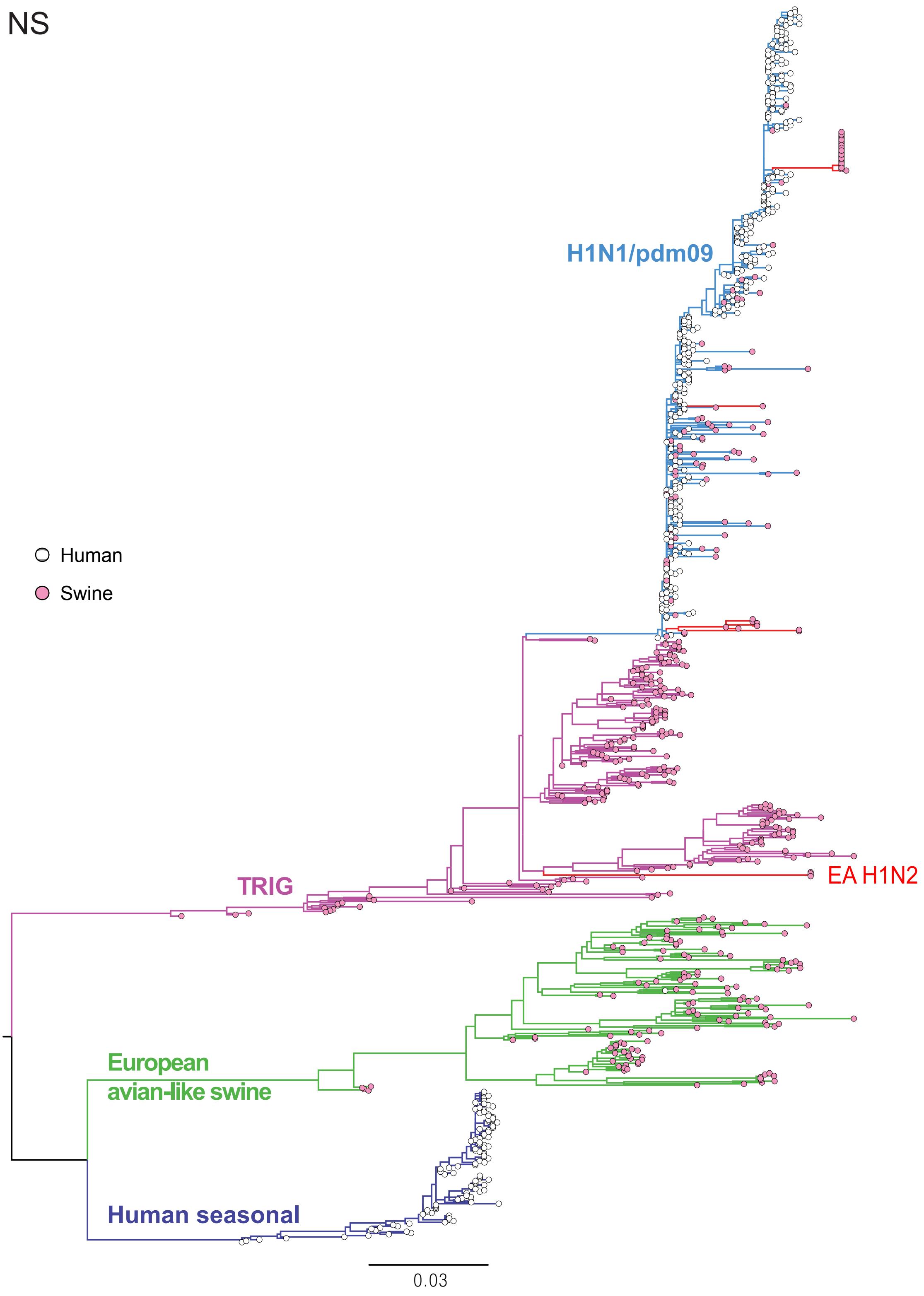


**Fig. S6**. Maximum likelihood phylogeny of the NS gene of human and swine influenza viruses. Coloured branches represent major lineages. Red branches denote swine sequences generated in this study. The novel EA H1N2 from pigs in Cambodia is indicated. Human and swine viruses are shown by pink and white tip circles. The scale bar represents the number of nucleotide substitutions per site.

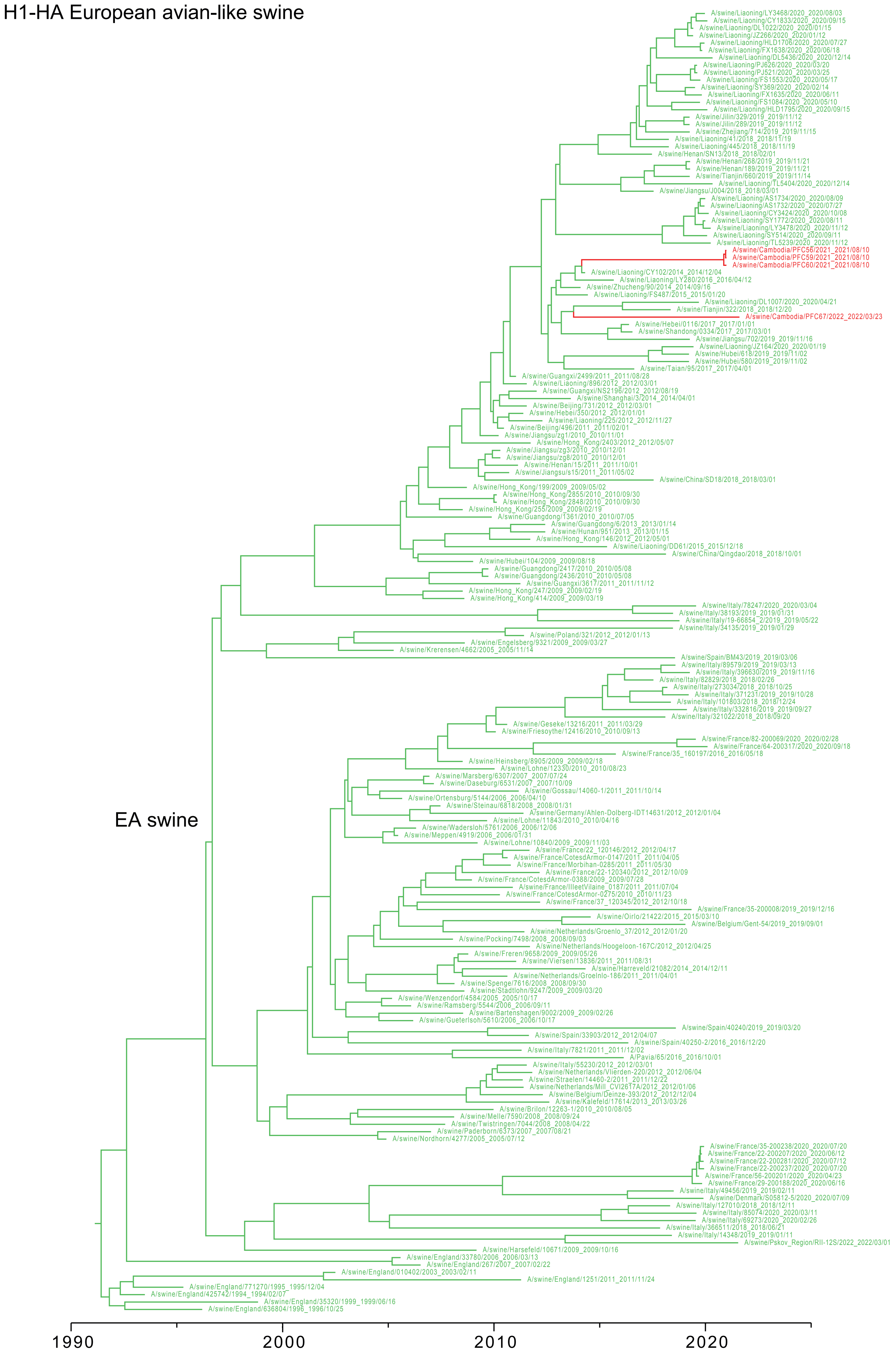


**Fig. S7.** Time scaled phylogeny of the H1-HA gene of European avian-like swine lineage. Red branches denote swine sequences generated in this study.

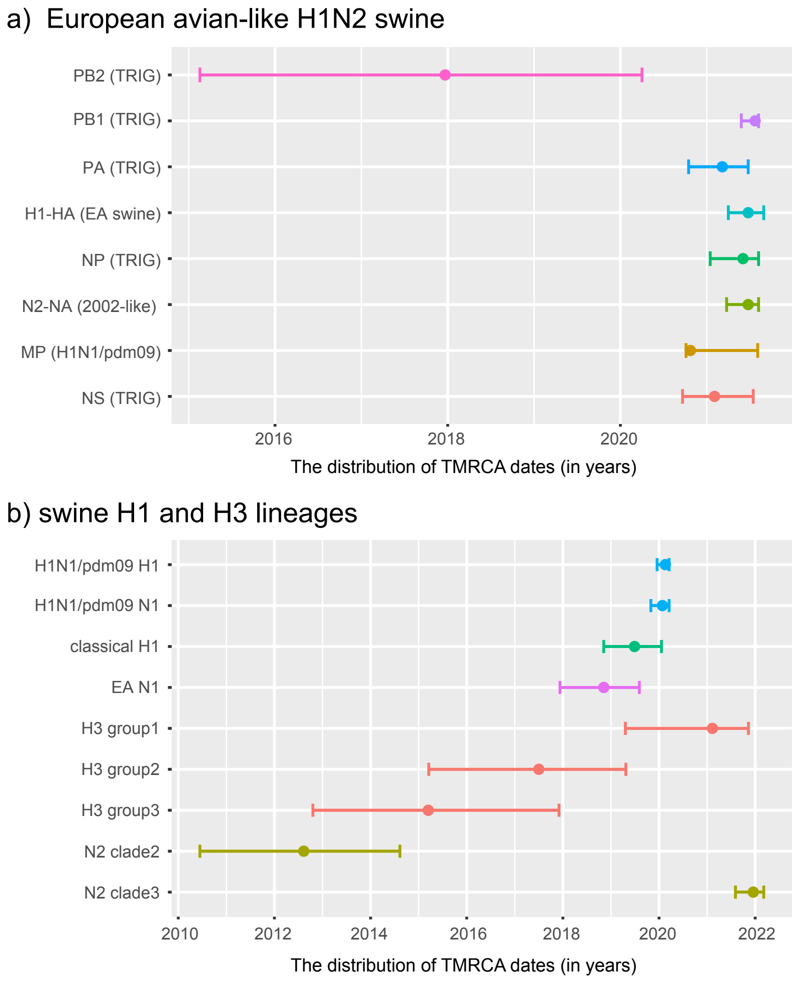


**Fig. S8.** The estimated time of the most recent common ancestors (TMRCAs) of different swIAV lineages from pigs in Cambodia. (a) The distribution of TMRCA dates of the Cambodian EA-swine lineage. Lineage name in brackets represent lineage designation. (b) The distribution of TMRCA dates of H1 and H3 lineages of Cambodia IAV viruses.

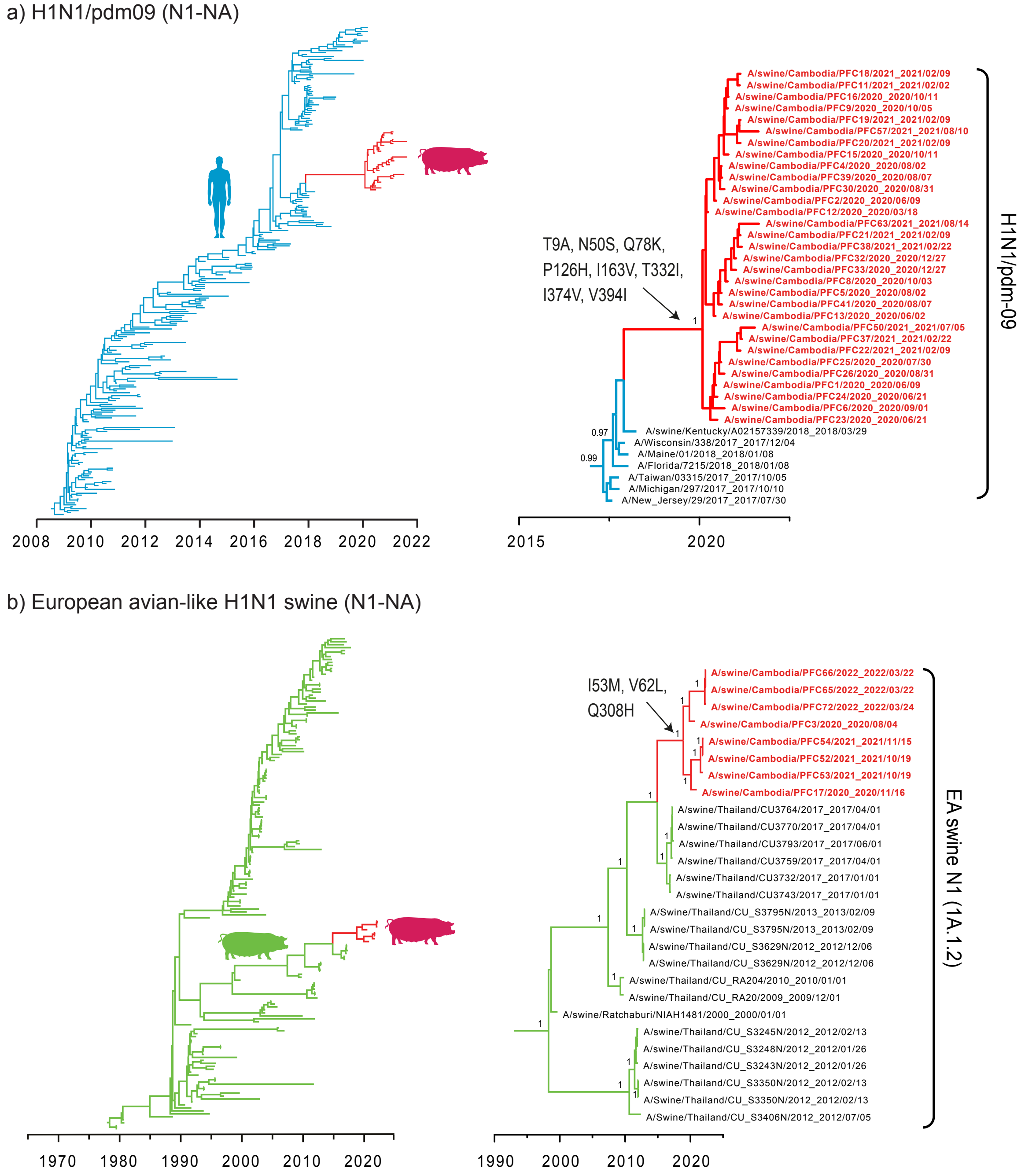


**Fig. S9.** The estimated time of the most recent common ancestors (TMRCAs) of different swIAV lineages from pigs in Cambodia. (a) The distribution of TMRCA dates of the Cambodian EA-swine lineage. Lineage name in brackets represent lineage designation. (b) The distribution of TMRCA dates of H1 and H3 lineages of Cambodia IAV viruses.

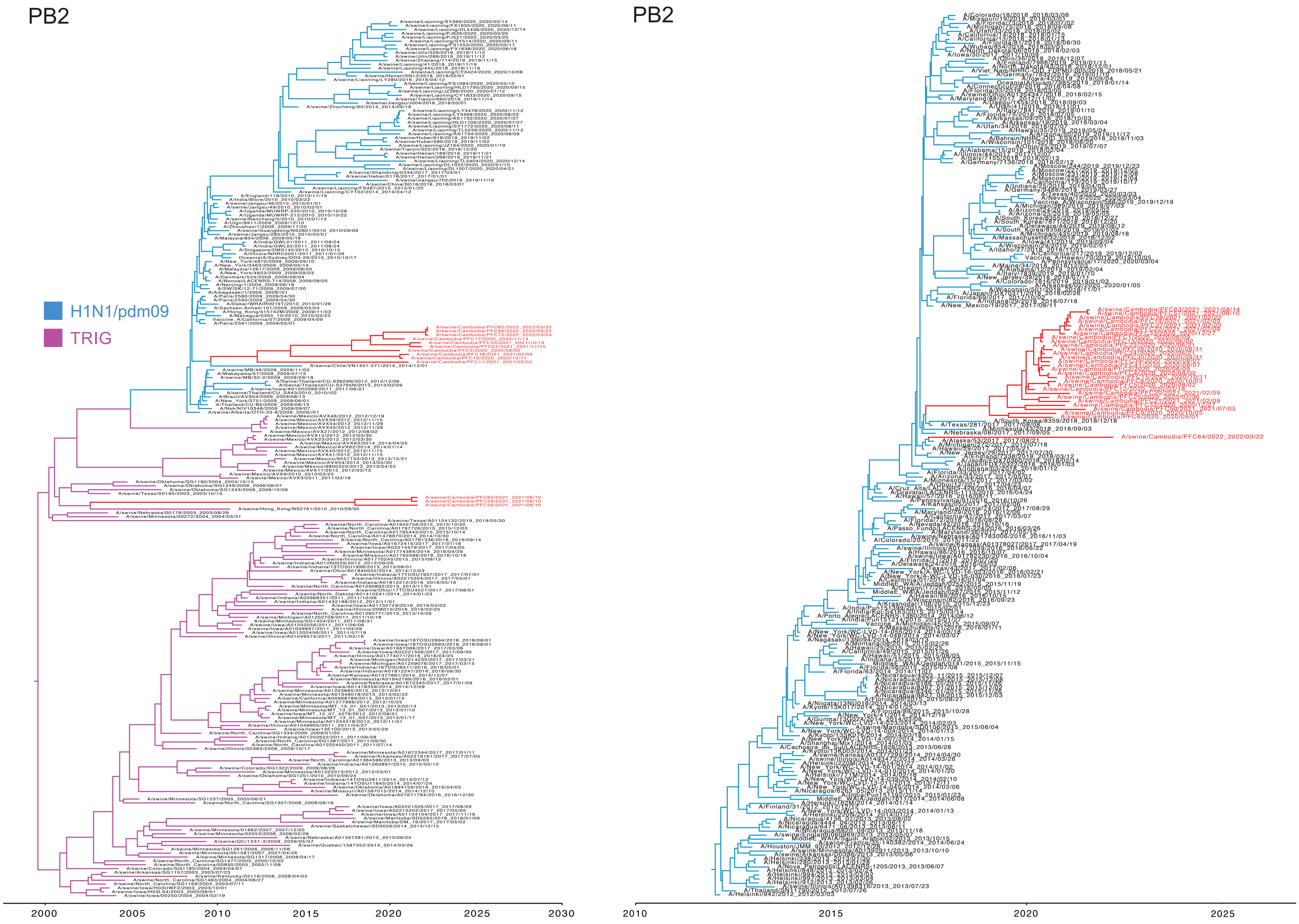


**Fig. S10.** Time scaled phylogenies of the PB2 gene of human and swine influenza viruses. The maximum clade credibility tree (MCC) on the right indicates swIAV sequences from Cambodian pigs are derived from H1N1/pdm-09 viruses, and their closely related viruses. The MCC on the left indicates the origins of swIAV sequences from Cambodian pigs and their closely related viruses. Red branches denote swine sequences generated in this study.

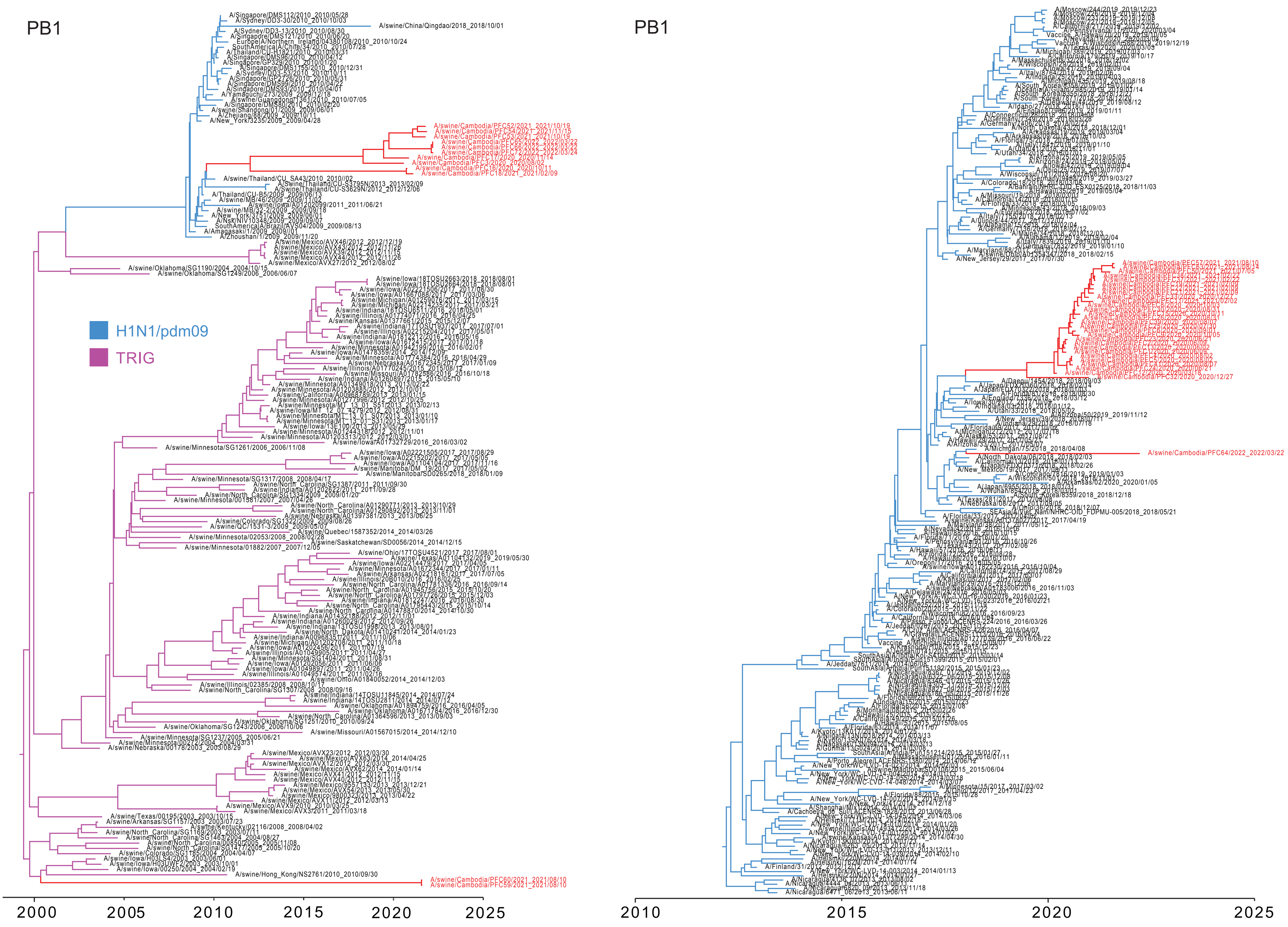


**Fig. S11.** Time scaled phylogenies of the PB1 gene of human and swine influenza viruses. The maximum clade credibility tree (MCC) on the right indicates swIAV sequences from Cambodian pigs are derived from H1N1/pdm-09 viruses, and their closely related viruses. The MCC on the left indicates the origins of swIAV sequences from Cambodian pigs and their closely related viruses. Red branches denote swine sequences generated in this study.

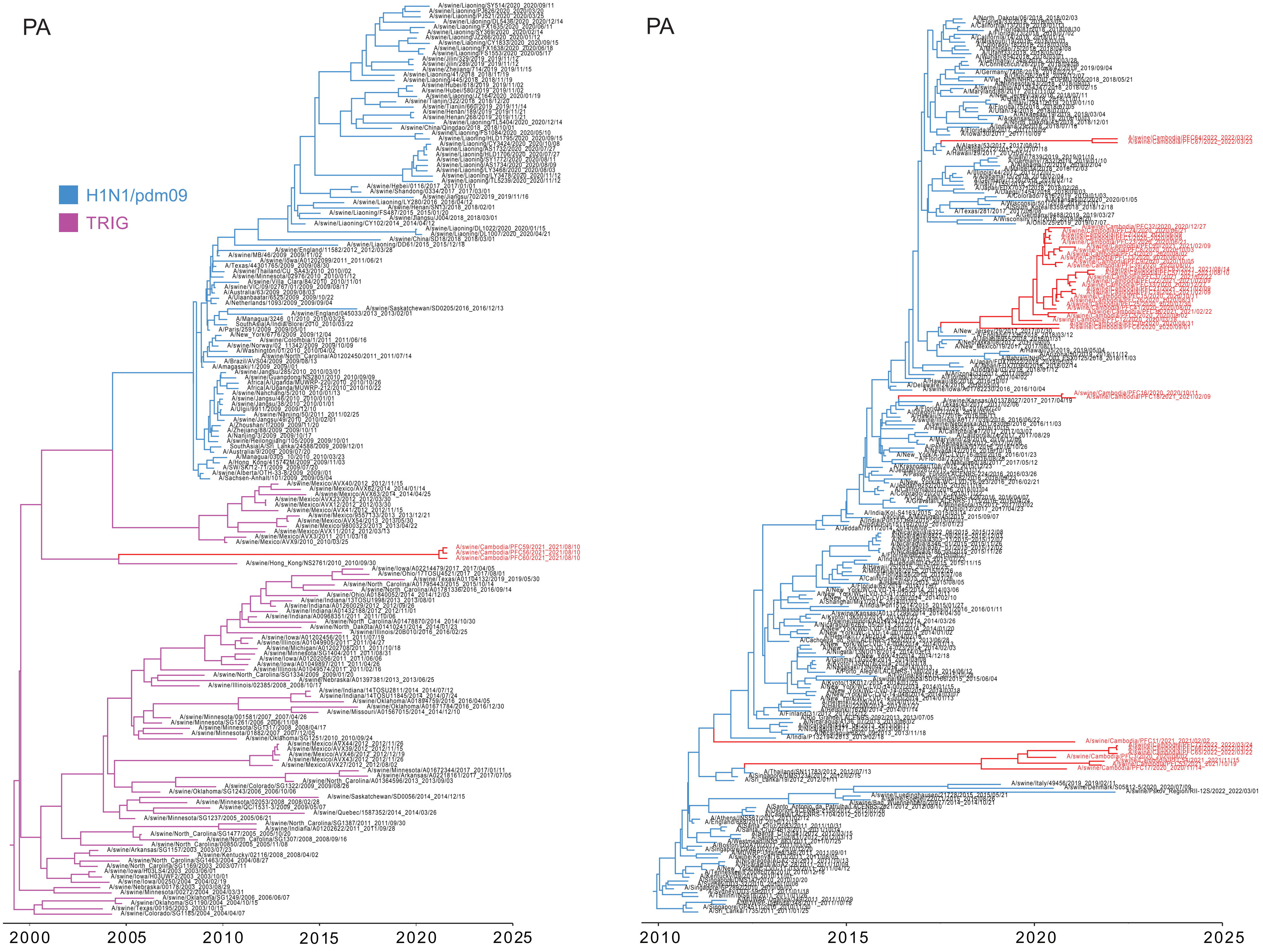


**Fig. S12.** Time scaled phylogenies of the PA gene of human and swine influenza viruses. The maximum clade credibility tree (MCC) on the right indicates swIAV sequences from Cambodian pigs are derived from H1N1/pdm-09 viruses, and their closely related viruses. The MCC on the left indicates the origins of swIAV sequences from Cambodian pigs and their closely related viruses. Red branches denote swine sequences generated in this study.

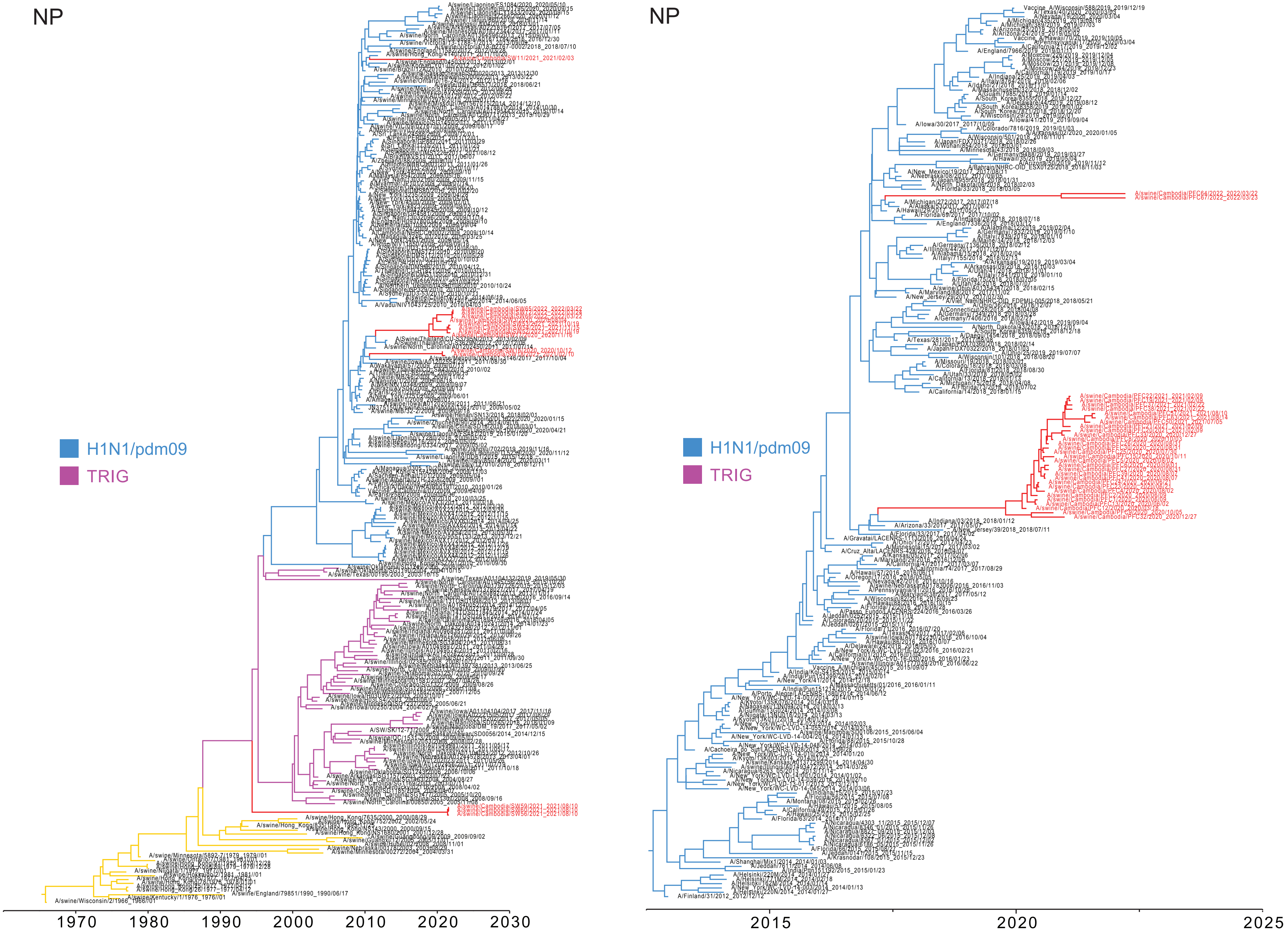


**Fig. S13.** Time scaled phylogenies of the NP gene of human and swine influenza viruses. The maximum clade credibility tree (MCC) on the right indicates swIAV sequences from Cambodian pigs are derived from H1N1/pdm-09 viruses, and their closely related viruses. The MCC on the left indicates the origins of swIAV sequences from Cambodian pigs and their closely related viruses. Red branches denote swine sequences generated in this study.

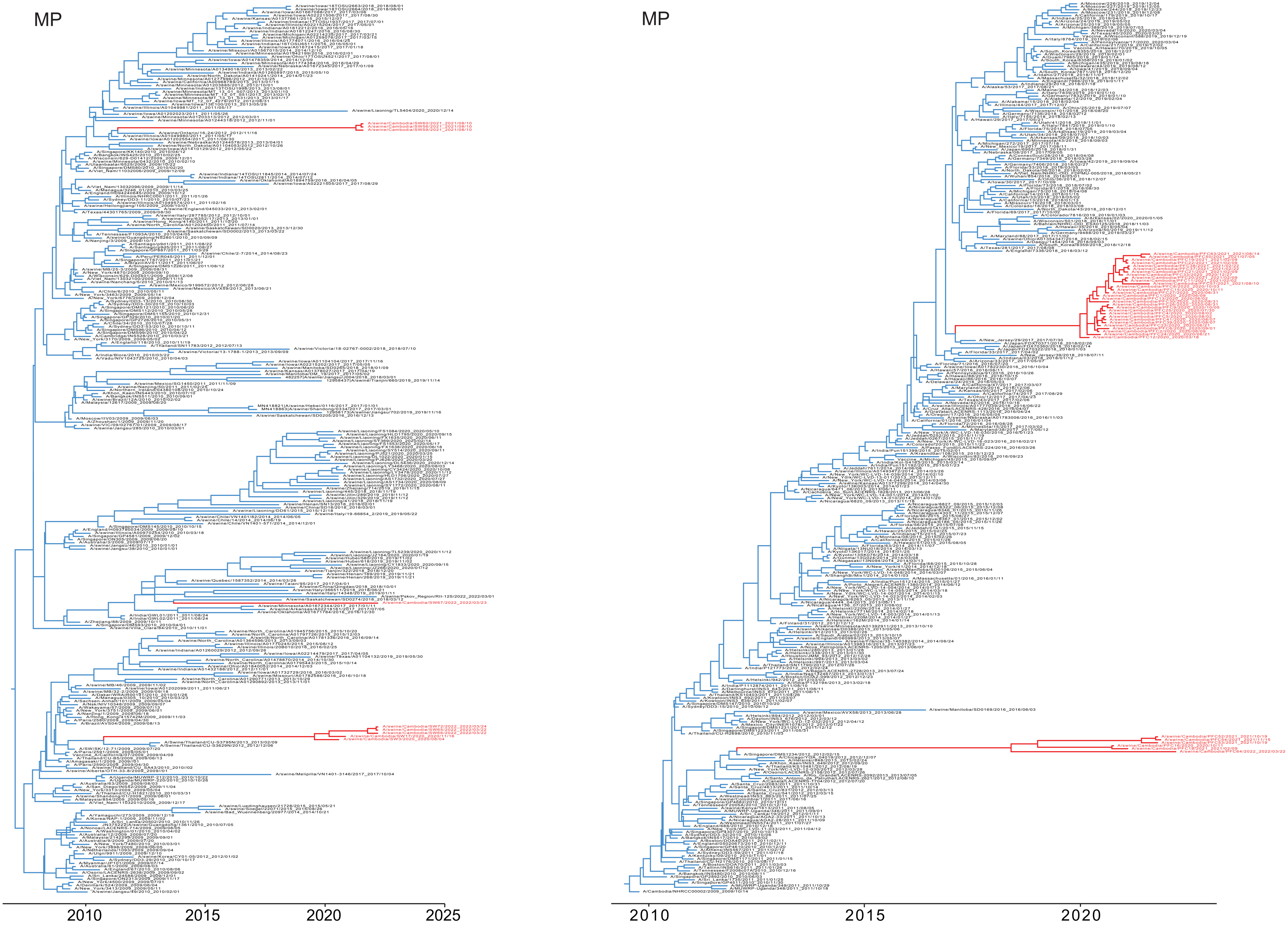


**Fig. S14.** Time scaled phylogenies of the MP gene of human and swine influenza viruses. The maximum clade credibility trees (MCC) indicate swIAV sequences from Cambodian pigs are derived from H1N1/pdm-09 viruses, and their closely related viruses. Red branches denote swine sequences generated in this study.

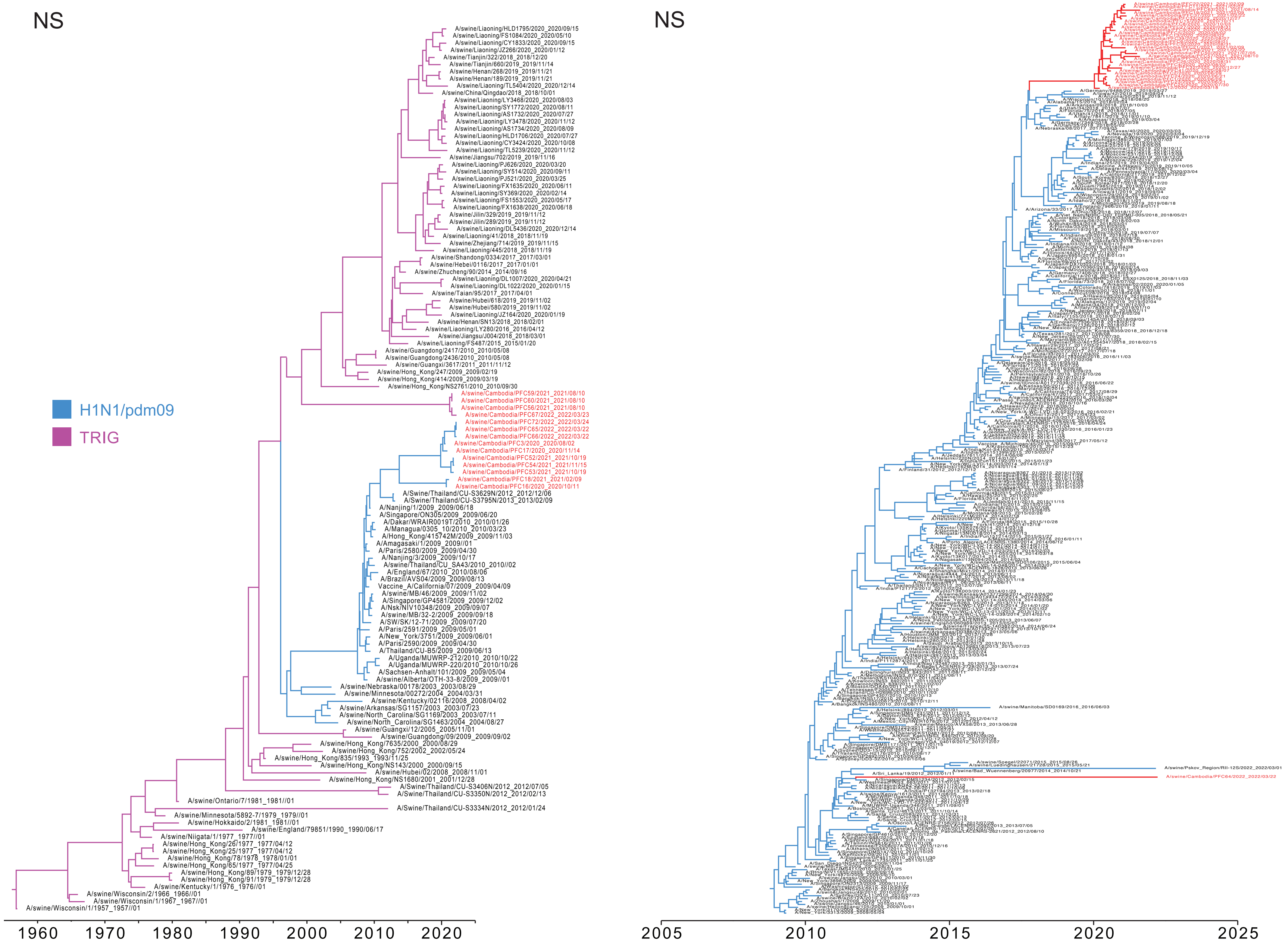


**Fig. S15.** Time scaled phylogenies of the NS gene of human and swine influenza viruses. The maximum clade credibility tree (MCC) on the right indicates swIAV sequences from Cambodian pigs are derived from H1N1/pdm-09 viruses, and their closely related viruses. The MCC on the left indicates the origins of swIAV sequences from Cambodian pigs and their closely related viruses. Red branches denote swine sequences generated in this study.

**Table S1.** Sequenced swine influenza A strains collected from pig slaughterhouses in Cambodia, 2020–2022.

| Strains | Province | District | Collection Date |
|---|---|---|---|
| A/swine/Cambodia/PFC1/2020 (H1N1) | Takeo | District 1 | 09/06/2020 |
| A/swine/Cambodia/PFC2/2020 (H1N1) | Takeo | District 1 | 09/06/2020 |
| A/swine/Cambodia/PFC3/2020 (H1N1) | Takeo | District 1 | 02/08/2020 |
| A/swine/Cambodia/PFC4/2020 (H1N1) | Takeo | District 1 | 02/08/2020 |
| A/swine/Cambodia/PFC5/2020 (H1N1) | Takeo | District 1 | 02/08/2020 |
| A/swine/Cambodia/PFC6/2020 (H1N1) | Takeo | District 1 | 01/09/2020 |
| A/swine/Cambodia/PFC8/2020 (H1N1) | Takeo | District 1 | 03/10/2020 |
| A/swine/Cambodia/PFC9/2020 (H1N1, H3N2) | Takeo | District 1 | 05/10/2020 |
| A/swine/Cambodia/PFC11/2021 (H1N1/H3) | Takeo | District 1 | 02/02/2021 |
| A/swine/Cambodia/PFC12/2020 (H1N1) | Kandal | District 9 | 18/03/2020 |
| A/swine/Cambodia/PFC13/2020 (H1N1) | Kandal | District 9 | 02/06/2020 |
| A/swine/Cambodia/PFC15/2020 (H1N1) | Kandal | District 8 | 11/10/2020 |
| A/swine/Cambodia/PFC16/2020 (H1N1, H3N2) | Kandal | District 8 | 11/10/2020 |
| A/swine/Cambodia/PFC17/2020 (H1N1) | Kandal | District 8 | 14/11/2020 |
| A/swine/Cambodia/PFC18/2021 (H1N1, H3N2) | Kandal | District 8 | 09/02/2021 |
| A/swine/Cambodia/PFC19/2021 (H1N1) | Kandal | District 8 | 09/02/2021 |
| A/swine/Cambodia/PFC20/2021 (H1N1/ H3) | Kandal | District 8 | 09/02/2021 |
| A/swine/Cambodia/PFC21/2021 (H1N1) | Kandal | District 8 | 09/02/2021 |
| A/swine/Cambodia/PFC22/2021 (H1N1, H3N2) | Kandal | District 8 | 09/02/2021 |
| A/swine/Cambodia/PFC23/2020 (H1N1) | Phnom Penh | District 5 | 21/06/2020 |
| A/swine/Cambodia/PFC24/2020 (H1N1) | Phnom Penh | District 5 | 21/06/2020 |
| A/swine/Cambodia/PFC25/2020 (H1N1) | Phnom Penh | District 7 | 30/07/2020 |
| A/swine/Cambodia/PFC26/2020 (H1N1/N2) | Phnom Penh | District 6 | 31/08/2020 |
| A/swine/Cambodia/PFC27/2020 (H1Nx) | Phnom Penh | District 6 | 31/08/2020 |
| A/swine/Cambodia/PFC30/2020 (H1N1, H3N2) | Phnom Penh | District 6 | 31/08/2020 |
| A/swine/Cambodia/PFC32/2020 (H1N1, H3N2) | Phnom Penh | District 7 | 27/12/2020 |
| A/swine/Cambodia/PFC33/2020 (H1N1) | Phnom Penh | District 7 | 27/12/2020 |
| A/swine/Cambodia/PFC37/2021 (H1N1) | Phnom Penh | District 6 | 22/02/2021 |
| A/swine/Cambodia/PFC38/2021 (H1N1/N2) | Phnom Penh | District 6 | 22/02/2021 |
| A/swine/Cambodia/PFC39/2020 (H1N1) | Kampong Speu | District 3 | 07/08/2020 |
| A/swine/Cambodia/PFC41/2020 (H1N1) | Kampong Speu | District 3 | 07/08/2020 |
| A/swine/Cambodia/PFC50/2021 (HxN1) | Phnom Penh | District 7 | 05/07/2021 |
| A/swine/Cambodia/PFC52/2021 (HxN1) | Takeo | District 1 | 19/10/2021 |
| A/swine/Cambodia/PFC53/2021 (H1N1) | Takeo | District 1 | 19/10/2021 |
| A/swine/Cambodia/PFC54/2021 (H1N1) | Takeo | District 1 | 15/11/2021 |
| A/swine/Cambodia/PFC56/2021 (H1N2) | Kandal | District 8 | 10/08/2021 |
| A/swine/Cambodia/PFC57/2021 (H1N1) | Kandal | District 8 | 10/08/2021 |
| A/swine/Cambodia/PFC59/2021 (H1N2) | Kandal | District 8 | 10/08/2021 |
| A/swine/Cambodia/PFC60/2021 (H1N2) | Kandal | District 8 | 10/08/2021 |
| A/swine/Cambodia/PFC63/2021 (H1N1) | Kandal | District 9 | 14/08/2021 |
| A/swine/Cambodia/PFC64/2022 (H3N2) | Kandal | District 9 | 22/03/2022 |

| | | | |
|---|---|---|---|
| A/swine/Cambodia/PFC65/2022 (H1N1) | Kandal | District 9 | 22/03/2022 |
| A/swine/Cambodia/PFC66/2022 (H1N1) | Kandal | District 9 | 22/03/2022 |
| A/swine/Cambodia/PFC67/2022 (H1N2/H3) | Kandal | District 9 | 23/03/2022 |
| A/swine/Cambodia/PFC72/2022 (H1N1/N2) | Kandal | District 8 | 24/03/2022 |

**Table S2.** Bayes factor of migration pathways of European avian-like H1 swine between geographic locations.

| From location | To destination | Bayes factor |
|---|---|---|
| East China | Europe | <3 |
| East China | North China | 22941.3 |
| East China | Northeast China | 22941.3 |
| East China | Southeast Asia | <3 |
| East China | South Central China | 22941.3 |
| East China | West China | 22941.3 |
| Europe | North China | <3 |
| Europe | Northeast China | <3 |
| Europe | Southeast Asia | 288.9 |
| Europe | South Central China | 9.2 |
| Europe | West China | <3 |
| Europe | East China | <3 |
| North China | Northeast China | 166 |
| North China | Southeast Asia | <3 |
| North China | South Central China | <3 |
| North China | West China | 7.3 |
| North China | East China | 4.5 |
| North China | Europe | <3 |
| Northeast China | Southeast Asia | 161 |
| Northeast China | South Central China | 5.9 |
| Northeast China | West China | <3 |
| Northeast China | East China | <3 |
| Northeast China | Europe | <3 |
| Northeast China | North China | 29.9 |
| South Central China | West China | 1037.8 |
| South Central China | East China | 22941.3 |
| South Central China | Europe | <3 |
| South Central China | North China | <3 |
| South Central China | Northeast China | 3819.2 |
| South Central China | Southeast Asia | <3 |
| Southeast Asia | South Central China | 8 |
| Southeast Asia | West China | <3 |
| Southeast Asia | East China | <3 |
| Southeast Asia | Europe | <3 |
| Southeast Asia | North China | <3 |
| Southeast Asia | Northeast China | <3 |
| West China | East China | <3 |
| West China | Europe | <3 |
| West China | North China | <3 |
| West China | Northeast China | <3 |
| West China | Southeast Asia | <3 |

| | | |
|---|---|---|
| West China | South Central China | <3 |

**Table S3.** Asymmetric diffusion rates between geographic locations for the European avian-like H1 lineage.

| | East China | Europe | North China | Northeast China | Southeast Asia | South Central China | West China |
|---|---|---|---|---|---|---|---|
| East China | - | 0.14 | 2.11 | 2.51 | 0.17 | 3.83 | 1.23 |
| Europe | 0.07 | - | 0.04 | 0.05 | 0.13 | 0.11 | 0.03 |
| North China | 0.62 | 0.23 | - | 0.93 | 0.24 | 0.57 | 0.38 |
| Northeast China | 0.30 | 0.26 | 0.53 | - | 0.38 | 0.50 | 0.31 |
| Southeast Asia | 0.24 | 0.21 | 0.22 | 0.21 | - | 0.48 | 0.23 |
| South Central China | 1.01 | 0.13 | 0.25 | 1.04 | 0.25 | - | 0.68 |
| West China | 0.19 | 0.19 | 0.20 | 0.21 | 0.20 | 0.23 | - |
